# A Leak K^+^ Channel TWK-40 Regulates Rhythmic Motor Program in *C. elegans*

**DOI:** 10.1101/2022.04.09.487752

**Authors:** Zhongpu Yue, Yi Li, Bin Yu, Yueqing Xu, Lili Chen, Jyothsna Chitturi, Jun Meng, Wesley Hung, Yuhang Tian, Sonia El Mouridi, Cuntai Zhang, Thomas Boulin, Mei Zhen, Shangbang Gao

**Affiliations:** Key Laboratory of Molecular Biophysics of the Ministry of Education, College of Life Science and Technology, Huazhong University of Science and Technology, Wuhan 430074, China; College of Biomedical Engineering, South-Central University for Nationalities, Wuhan 430074, China; Department of Geriatrics, Tongji Hospital of Tongji Medical College, Huazhong University of Science and Technology, Wuhan, China; Lunenfeld-Tanenbaum Research Institute, Mount Sinai Hospital, University of Toronto, Toronto, Ontario M5G 1X5, Canada; Univ Lyon, Université Claude Bernard Lyon 1, MeLiS, CNRS UMR 5284, INSERM U1314, Institut NeuroMyoGène, Lyon 69008, France; King Abdullah University of Science and Technology (KAUST), Biological and Environmental Sciences and Engineering Division (BESE), Thuwal 23955–6900, Kingdom of Saudi Arabia

**Keywords:** *twk-40*, K_2P_ channel, motor rhythm, defecation, Ca^2+^ oscillation, resting membrane potential

## Abstract

Leak potassium (K^+^) currents, conducted by two-pore domain K^+^ (K_2P_) channels, are critical for the stabilization of the membrane potential. The effect of K_2P_ channels on motor rhythm remains enigmatic. We show here that the K_2P_ TWK-40 regulates the rhythmic defecation motor program (DMP) in *Caenorhabditis elegans*. Disrupting TWK-40 suppresses the expulsion defects of *nlp-40* and *aex-2* mutants. By contrast, a gain-of-function (*gf*) mutant of *twk-40* significantly reduces the expulsion frequency per DMP cycle. *In situ* whole-cell patch clamping demonstrates that TWK-40 forms an outward current that hyperpolarize the resting membrane potential of DVB neuron. In addition, TWK-40 substantially contributes to the rhythmic activity of DVB. Specifically, DVB Ca^2+^ oscillations exhibit obvious defects in *twk-40* mutants. Expression of TWK-40(*gf*) in DVB recapitulates the expulsion deficiency of the *twk-40(gf)* mutant, and inhibits DVB Ca^2+^ oscillations in both wild-type and *twk-40(lf)* animals. Moreover, DVB innervated enteric muscles also exhibit rhythmic Ca^2+^ defects. Taken together, these results demonstrate that TWK-40 is an essential neuronal inhibitor of DMP, thus revealing a cellular mechanism linking leak K_2P_ channels with rhythmic motor activity.

## Introduction

Two-pore domain potassium (K_2P_) channels conduct K^+^ leak currents. In contrast to voltage-gated K^+^ channels, K_2P_ channels are mostly voltage-independent and non-inactivating channels, which stabilize the cell’s resting membrane potential (Enyedi and Czirjak, 2010). Along with most K_2P_ channels profiling by heterologous expression systems or in cultured primary cells, the channel associated behaviors and the intrinsic physiological characterization of specific K_2P_ channel have remained largely unknown.

The human genome encodes 15 K_2P_ channels, which are grouped into six families based on their functional resemblance and structural similarity (Goldstein et al., 2005; Lesage and Lazdunski, 2000). Aberrant functions of K_2P_ channels have been implicated in various disorders associated with genetic variation. For example, mutations of human TASK-3 (KCNK9 p.Gly236Arg) cause *KCNK9* Imprinting Syndrome, a pediatric neurodevelopmental disease with severe feeding difficulties, delayed development and intellectual disability (Graham et al., 2016). With predicted conservation of amino acid sequences across species, K_2P_ channels associated genetic and functional investigations in animal models are emerging (Buckingham et al., 2005; Lalevee et al., 2006). For instance, cardiac-specific inactivation or overexpression of Ork1, a *Drosophila* two-pore domain potassium channel, led to an increase or a complete arrest of fly heart beating, respectively (Lalevee et al., 2006).

The nematode *Caenorhabditis elegans* (or *C. elegans*) genome contains a large family with at least 47 K_2P_ genes (Bargmann, 1998; Buckingham *et al*., 2005). This large set of K_2P_ channels may allow for exceptionally fine ‘tuning’ of the firing activity of individual cells within the very compact nematode nervous system (Salkoff et al., 2001; Witvliet et al., 2021). Encouraged by the increasing understanding of the molecular and cellular wiring of the worm’s neural network (Bargmann, 1998; Cook et al., 2019; White et al., 1986; Witvliet *et al*., 2021), we use *C. elegans* to explore the functional complexity of K_2P_ channels as potential regulators of rhythmic motor behaviors *in vivo* (Ben Soussia et al., 2019; Wang et al., 1999).

The *C. elegans* defecation motor program (DMP) exhibits a highly coordinated ultradian rhythm. It is achieved by periodically activating a stereotyped sequence of muscle contractions, including the initial contraction of posterior body wall muscles (pBoc), followed by the second contraction of anterior body wall muscles (aBoc), and the final contraction of enteric muscles (EMC) (Thomas, 1990). Driven by the synchronizing activity of electrically-coupled DVB and AVL motor neurons (Choi et al., 2021), EMC leads to the robust expulsion (Exp) of gut contents (**Figure 1A**) (Dal Santo et al., 1999; Liu and Thomas, 1994). Unlike most rhythmic motor circuits that are composed of a central pattern generator (CPG) neural network or self-oscillating pacemaker neurons (Marder and Calabrese, 1996), the expulsion rhythm is precisely timed by endogenous calcium oscillation signals from the intestine (Branicky and Hekimi, 2006; McIntire et al., 1993a). The intestinal IP_3_ receptor/*itr-1*-driven pacemaker Ca^2+^ oscillations trigger a Ca^2+^-dependent secretion of neuropeptide NLP-40, which activates the neuronal G protein-coupled receptor (GPCR) AEX-2 in DVB and AVL (Mahoney et al., 2008; Nehrke et al., 2008; Teramoto and Iwasaki, 2006; Wang et al., 2013). GABA release from DVB (McIntire et al., 1993b), cyclically initiated by NLP-40 (Wang *et al*., 2013), activates the muscular excitatory GABA-gated cation channel EXP-1 to drive the final contraction of enteric muscles (**Figure 1B**) (Beg and Jorgensen, 2003). Deficiency of either NLP-40, AEX-2 or EXP-1 results in serious expulsion defects in *C. elegans*. Although the intestinal Ca^2+^ oscillation sets the instructive timing signal of the expulsion rhythm (Branicky and Hekimi, 2006; McIntire *et al*., 1993a), the cellular processes that control the expulsion rhythm in the intestine-neuron-muscle circuit are still unclear.

**Figure 1.**
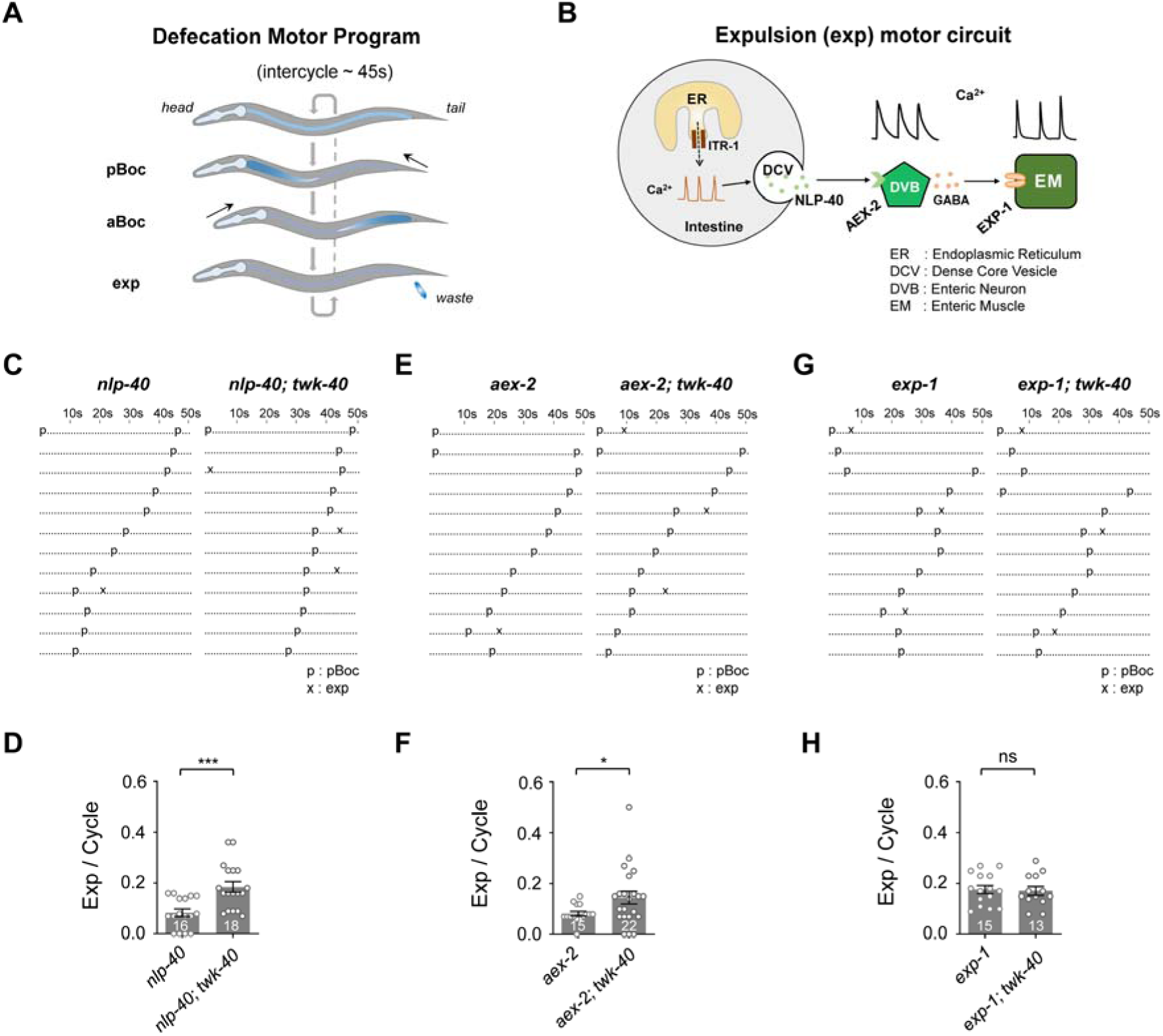
Loss of function of twk-40 suppresses expulsion defects in nlp-40 and aex-2 mutants. (**A**) Schematic representation of the <u>d</u>efecation <u>m</u>otor <u>p</u>rogram (DMP), which consists of three distinct sets of muscle contractions: the posterior body muscle contraction (pBoc), the anterior body muscle contraction (aBoc), and the expulsion muscle contraction (exp or EMC). The regular defecation cycle period is approximately 45 sec. (**B**) Schematic representation of the expulsion motor circuit, including the intestine, DVB neuron and enteric muscles (EM). ER, endoplasmic reticulum; ITR-1, inositol 1,4,5-trisphosphate receptor; DCV: dense core vesicle. (**C, E, G**) Ethograms of defecation behavior in *nlp-40* and *nlp-40; twk-40* double mutants, *aex-2* and *aex-2; twk-40* double mutants, and *exp-1* and *exp-1; twk-40* double mutants, respectively. Each dot and character represent 1 s. Letters ‘p’ and ‘x’ represent pBoc and exp, respectively. (**D, F, H**) Quantifications of the expulsion frequency per defecation cycle. The expulsion deficiency was partially recovered by loss of *twk-40* in *nlp-40* and *aex-2* mutants, but not in *exp-1* mutants. All data are expressed as means ± SEM. Student’s *t*-test was used. Statistical significance is indicated as follows: ns, not significant, \**P* < 0.05, \*\*\**P* < 0.001 in comparison with that as denoted.

We show here that loss of function of *twk-40* – a two-pore domain potassium channel subunit – specifically suppresses the expulsion defects of *nlp-40* and *aex-2* mutants, but not of *exp-1* mutant. TWK-40 exhibits the common structural features of K_2P_ channel based on sequence analyses. The single subunit of TWK-40 has four transmembrane regions and two pore-forming domains (**Figure 2A**), which exhibit sequence similarity with the mammalian TASK (tandem-pore-acid-sensing K^+^) subfamily (**Supplementary Figure 1A**). A gain of function mutant of *twk-40(bln336)*, generated by CRISPR/Cas9 gene editing (Ben Soussia *et al*., 2019), induces an expulsion defect by reducing the frequency of expulsion per DMP cycle. Moreover, *in-situ* whole-cell patch clamping revealed a *twk-40*-dependent outward K^+^ current that regulates the resting membrane potential (RMP) in DVB neurons, in which the RMP was depolarized in *twk-40(lf)* and hyperpolarized in *twk-40(gf)* mutant, respectively. In combination with real-time Ca^2+^ imaging, we provide several lines of evidence to demonstrate that TWK-40 regulates rhythmic motor behaviors – including defecation – by cell-autonomously regulating DVB’s resting membrane potential.

**Figure 2.**
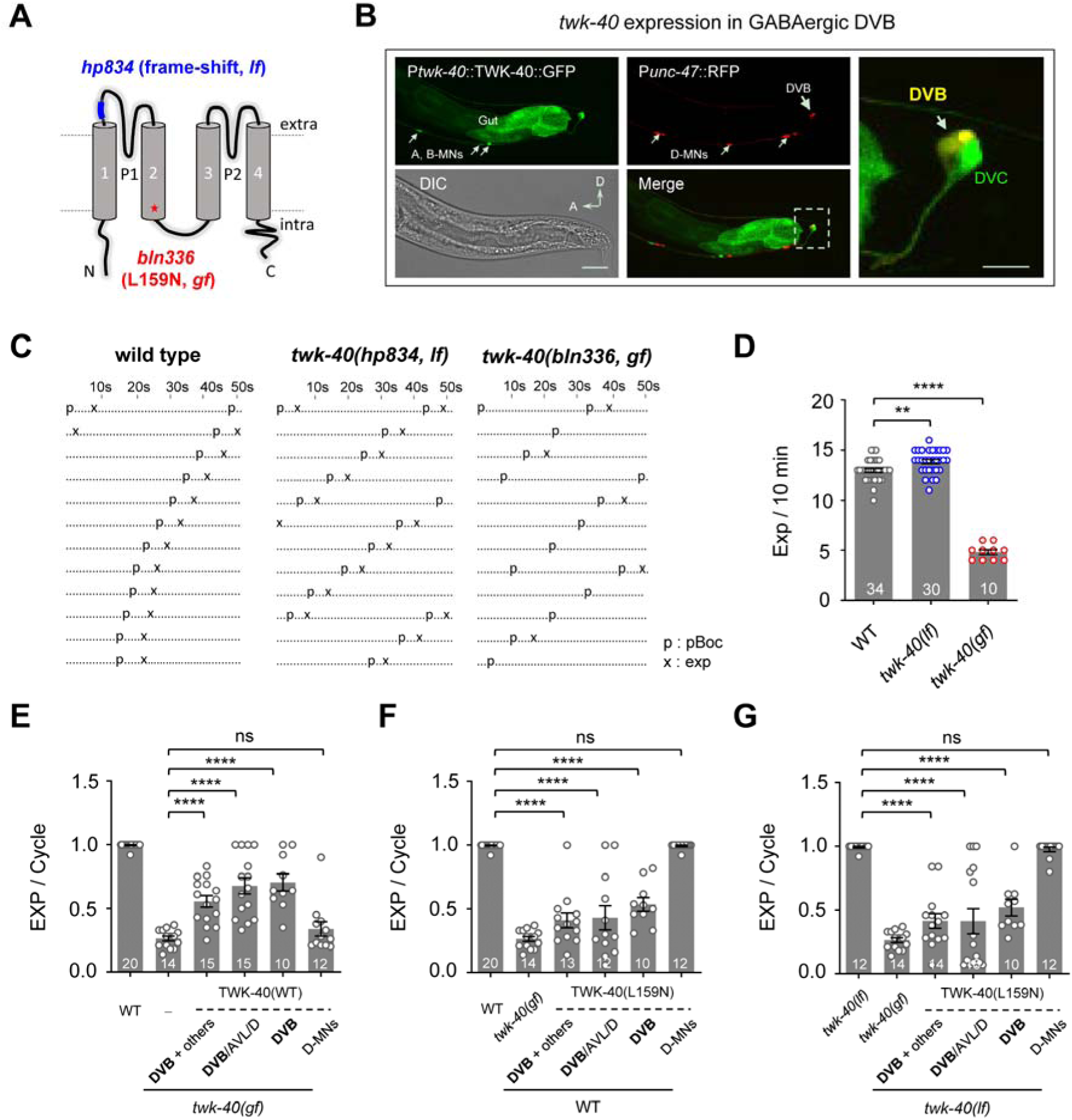
twk-40 regulates expulsion rhythm from DVB neuron. (**A**) Diagram of the TWK-40 K^+^ channel and the mutation loci. P1 and P2, pore domains; 1-4 transmembrane domains; *fs*, predicted frameshift mutation; *gf*, gain-of-function mutation. (**B**) *twk-40* is expressed in DVB based on co-localization of P*twk-40*::TWK-40::sl2dGFP and P*unc-47*::RFP. GFP was observed in the ventral excitatory motor neurons, a few head and tail neurons, and the intestine. Scale bar, 20 μm. *Right*, zoomed view of *twk-40* expression in DVB and DVC. Scale bar, 5 μm. (**C**) Ethograms of defecation behavior in wild type, *twk-40(bln336, gf)* and *twk-40(hp834, lf)* animals. Each dot and character represent 1 s. Letters ‘p’ and ‘x’ represent pBoc and exp, respectively. (**D**) Quantifications of the expulsion frequency per defecation cycle. Compared to wild-type animals, the expulsion deficiency was significantly reduced in *twk-40(bln336, gf)*, but not in *twk-40(hp834, lf)*. (**E**) Neuronal expression of wild-type TWK-40 (TWK-40(WT)) in DVB neurons rescues the expulsion deficiency of *twk-40(bln336, gf)* mutants. (**F, G**) Neuronal expression of gain-of-function TWK-40(L159N) in DVB in wild type (F) or *twk-40(hp834)* (G) recapitulates the expulsion deficiency of *twk-40(gf)* mutant. The number of tested animals is indicated for each strain. All data are expressed as means ± SEM. One-way ANOVA test was used, in which: ns, not significant, \*\**P* < 0.01, \*\*\*\**P* < 0.0001 relative to wild type or as indicated.

Remarkably, we also found recently that TWK-40 modulates global locomotor activity by regulating the activity of the AVA premotor interneuron (Meng et al., 2022). Briefly, *twk-40(lf)* mutant animals exhibit exaggerated body bends during both forward and backward movement, and more frequent and prolonged reversals. *twk-40(gf)* mutant animals, by contrast, exhibit strongly altered locomotion including obvious decreased body bending and velocity in both forward and backward undulation. Heterologous expression of the TWK-40 in HEK293T cells produced a voltage-insensitive K^+^ selective K_2P_-like ion channel. Therefore, the K_2P_ channel TWK-40 is required for the generation of multiple motor rhythms, including the defecation and locomotion motor programs.

## Results

### twk-40(lf) suppresses the expulsion defect of nlp-40 and aex-2 mutants

To determine a potential role of the TWK-40 K_2P_ channel in the cellular mechanisms controlling rhythmic DMP activity, we took advantage of three *exp* mutants that disrupt DMP in different tissues within the expulsion motor circuit (**Figure 1B**). First, *nlp-40(tm4085)*, an instructive neuropeptide from the intestinal pacemaker that delivers temporal information to the DVB neuron (Wang *et al*., 2013). Second, *aex-2(sa3)*, a G-protein-coupled receptor that functions as the DVB receptor for NLP-40 (Mahoney *et al*., 2008; Wang *et al*., 2013). And the third, *exp-1(sa6)*, which encodes an excitatory GABA receptor expressed in enteric muscles (Beg and Jorgensen, 2003; Lee et al., 2005). Loss-of-function mutants of these genes induced severe expulsion defects (**Figure 1C, E, G**) by disrupting the normal function of the intestine, DVB neuron and enteric muscles, respectively.

We reasoned that a loss of the potassium conductance *twk-40* would increase the excitability of cells within the circuit and compensate the inhibitory effect of the *nlp-40*, *aex-2*, or *exp-1* mutations. Hence, we combined the *twk-40* allele *hp834* (**Figure 2A**) (Meng *et al*., 2022) with each DMP mutation. *hp834* contains a 7 bp deletion of the sequence (GTTCGAG) at the base of 127-133 exon before the first pore-forming domain of T28A8.1a (**Figure 2A**), causing a frameshift and subsequent premature stop codon of *twk-40a* (Meng *et al*., 2022), one of three annotated isoforms of *twk-40* (www.wormbase.org). We found that *twk-40(hp834)* significantly improved the expulsion defect in *nlp-40* and *aex-2* mutants (**Figure 1C-F**). Specifically, the expulsion frequency per DMP was increased in *nlp-40(tm4085); twk-40(hp834)* (0.18±0.02 Exp per cycle) or *aex-2(sa3); twk-40(hp834)* (0.15±0.02 Exp per cycle), when compared to the single mutants of *nlp-40(tm4085)* (0.08±0.01 Exp per cycle) and *aex-2(sa3)* (0.08±0.01 Exp per cycle), respectively (**Figure 1C-F**). Similar expulsion defect suppression was observed in another *twk-40(lf)* allele *hp733* (**Supplementary Figure 1B**), which harbors a missense mutation that changes an amino acid at the end of the first transmembrane domain (E58K) (Meng *et al*., 2022). By contrast, no substantial suppression of the expulsion defect of *exp-1* mutants was observed for either loss-of-function allele of *twk-40*. Indeed, no difference in expulsion frequency was observed between *exp-1(sa6)* (0.17±0.01 Exp per cycle) and *exp-1(sa6); twk-40(hp834)* (0.17±0.01 Exp per cycle) or *exp-1(sa6); twk-40(hp733)* (0.16±0.01 Exp per cycle) double mutants, respectively (**Figure 1G, H**, **Supplementary Figure 1B**). These results demonstrated that *twk-40* is selectively required for rhythmic defecation regulation, upstream of *exp-1*.

### A twk-40 gain-of-function mutation reduces expulsion frequency

The selective suppression of *exp* defects in distinct *exp* mutants suggested that *twk-40, per se,* may regulate the DMP. Indeed, in *twk-40(hp834, lf)* single mutants, the expulsion number was modestly but significantly increased (13.87±0.21/10 min) compared to wild-type animals (12.97±0.19/10 min) (**Figure 2C, D**). Similar results were also observed in *twk-40(hp733)* mutants. These results confirm that *twk-40* regulates the DMP negatively.

To further confirm the involvement of *twk-40* in the defecation motor program, we examined the expulsion step in a gain-of-function mutant allele of *twk-40, bln336* (**Figure 2A**). These mutants harbor a pan-K_2P_ activating mutation (Ben Soussia *et al*., 2019) that promotes the gating of vertebrate and invertebrate K_2P_ channels. Indeed, heterologous expression of the corresponding TWK-40(L159N) channel in HEK293T cells showed ∼5 fold current increase from the wild-type TWK-40 (Meng *et al*., 2022). Consistent with an inhibitory function of *twk-40*, we observed a strongly reduced expulsion frequency (4.8±0.25/10 min) in *twk-40(bln336, L159N*) animals, i.e. approximately 25% of wild type (12.97±0.19/10 min) (**Figure 2C, D**). Thus, gain-of-function of *twk-40* inhibits the expulsion rhythm, which encouraged us to further investigate the functional effects of *twk-40* loss-and gain-of-function mutants.

### twk-40 regulates expulsion in the DVB neuron

To pinpoint the exact role of *twk-40* in the DMP, we first assessed its cellular focus of action. Loss of *twk-40* suppressed the *exp* defect of *nlp-40* and *aex-2* mutants but not of *exp-1*, indicating that *twk-40* regulates the expulsion frequency from the intestine and/or expulsion neurons. Indeed, functional TWK-40 driven by a small fragment of its upstream region (P*twk-40*::TWK-40::sl2dGFP, Methods) revealed the expression exclusively in the nervous system and intestine (**Figure 2B**). The neuronal expression pattern includes the excitatory ventral motor neurons (A-and B-types), but no expression was observed in inhibitory D-motor neurons. Meanwhile, sparse labeling of head and tail neurons was also observed, including the DVC interneuron (strong) and the excitatory GABAergic DVB motor neuron/interneuron (moderate) (**Figure 2B**).

We then systematically tested the tissue or cell requirements of *twk-40* for the expulsion behavior. A wild-type TWK-40 cDNA driven by different promoters was competitively expressed in *twk-40(bln336, gf)* mutant (0.26±0.02 Exp per cycle), in an attempt to reduce its severe *exp* defect. Expression of wild-type TWK-40 (i.e., TWK-40(WT)) by the short promoter P*twk-40* (Methods) rescued the animal’s expulsion frequency to (0.55±0.04 Exp per cycle) (**Figure 2E**). Expression of TWK-40(WT) in GABAergic DVB/AVL/D-motor neurons (P*unc-47*), also significantly restored the DMP (0.67±0.06 Exp per cycle). More importantly, when we restored TWK-40(WT) expression exclusively in the DVB neuron (P*flp-10*)(Choi *et al*., 2021), the expulsion defect of *twk-40(bln336, gf)* mutant was rescued (0.7±0.06 Exp per cycle). In contrast, expression of the TWK-40(WT) in the intestine (P*ges-1*, 33.2±1.9%) (**Supplementary Figure S1C**) or D-motor neurons (P*unc-25*s, 33.8±5.6%) (Jin et al., 1999), however, did not restore a normal expulsion frequency (**Figure 2E**). Therefore, consistent with the expression pattern, these results suggest that regulation of the expulsion rhythm by *twk-40* requires DVB neurons.

To further confirm this notion, we ectopically expressed the TWK-40(L159N) gain-of-function mutant in DVB neurons. Indeed, in the wild-type N2 background, TWK-40(L159N) expression in DVB (P*twk-40*, P*unc-47* or P*flp-10*) resembled the expulsion defect as found in *twk-40(bln336)* mutant animals, exhibiting strongly reduced expulsion frequency per cycle (**Figure 2F**). TWK-40(L159N) expression only in D-MNs (P*unc-25*s), however, could not replicate this defect. Similar phenotypes were also observed by expressing TWK-40(L159N) in a *twk-40(lf)* mutant context (**Figure 2G**).

Taken together, these results demonstrate that *twk-40* is sufficient and necessary in DVB neurons for the regulation of the expulsion rhythm.

### Rhythmic Ca^2+^ oscillation of the DVB neuron is disrupted in twk-40 mutants

To determine whether *twk-40* directly regulates DVB’s neuronal activity, we performed Ca^2+^ imaging in animals expressing the genetically encoded Ca^2+^ sensor GCaMP6s in the DVB neuron (**Figure 3A**) (Li et al., 2023). We conducted the Ca^2+^ imaging in a liquid environment on restricted animals, in which DVB exhibited tight correlation of rhythmic Ca^2+^ oscillations with cyclic expulsion behavior (**Figure 3B-D**). In wild-type animals, DVB fired periodic Ca^2+^ spikes (or Ca^2+^ oscillations) with a frequency of 4.5±0.3 Hz/300s (**Figure 3B, E, F**). To verify these Ca^2+^ spikes represent pacemaker-activated DVB activation, we examined their dependence on NLP-40, the instructive neuropeptide from the intestinal pacemaker that delivers temporal information to the DVB neuron (Wang *et al*., 2013). Consistent with previous reports (Choi *et al*., 2021; Wang *et al*., 2013), we observed severe decrease in both DVB Ca^2+^ and expulsion frequency in the *nlp-40(tm4085)* mutant (**Supplementary Figure S2A-C**). These results demonstrate that the activation dynamics of DVB are indeed dependent on the NLP-40-initiated pacemaker signal.

**Figure 3.**
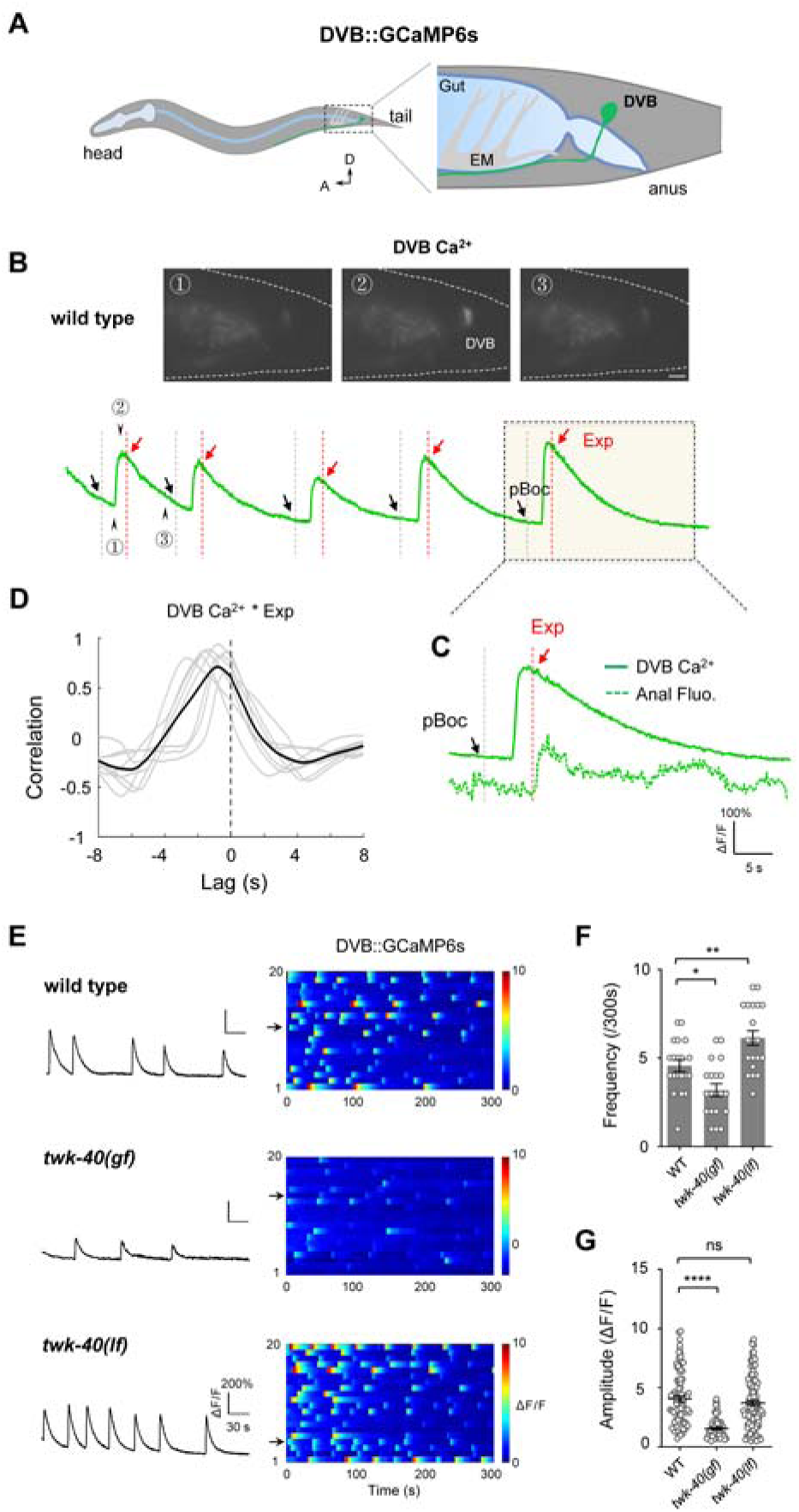
twk-40 inhibits DVB Ca^2+^ oscillation. (**A**) Schematic representation of *C. elegans* DVB neuron and enteric muscle (EM) in a lateral view. The genetically-encoded Ca^2+^ indicator GCaMP6s was expressed in DVB to measure its neuronal activity. (**B**) Representative spontaneous Ca^2+^ oscillation imaged by GCaMP in DVB neuron. *Upper*, from left to right, Ca^2+^ signal of DVB in (1) the quiescent state, (2) at peak Ca^2+^ signal and (3) upon return to the basal state. Scale bar, 5 μm. *Bottom*, representative recording of Ca^2+^ oscillations in DVB over time. Arrow heads indicate the corresponding time points in upper panels. (**C**) Simultaneous recording of DVB Ca^2+^ activity and expulsion action that observed by anal bacterial fluorescence. (**D**) Cross-correlation between DVB Ca^2+^ and Exp. Faint lines indicate the results from individual Ca^2+^ transient, and the black line indicates mean value. Black dashed line denotes tight correlation between DVB Ca^2+^ and Exp action with a ∼ 0.8 s delay. (**E**) Representative Ca^2+^ activity (*left*) and color maps (*right*) of DVB neurons in wild type, *twk-40(bln336, gf)* and *twk-40(hp834, lf)* mutants, respectively. (**F, G**) Significant reduction of Ca^2+^ transient frequency and amplitude in *twk-40(bln336, gf)* mutant, n=20. Reduction of frequency but not amplitude in *twk-40(hp834, lf)* mutants. All data are expressed as means ± SEM. One-way ANOVA test was used, in which: ns, not significant, \**P* < 0.05, \*\**P* < 0.01, \*\*\*\**P* < 0.0001 in comparison with that as denoted.

In *twk-40(bln336, gf)* mutants, the frequency of Ca^2+^ spikes was significantly reduced to 3.2±0.3 Hz/300s (**Figure 3E, F**). Interestingly, the Ca^2+^ amplitude was also diminished in *twk-40(bln336, gf)* mutants (**Figure 3G**). Conversely, in *twk-40(hp834, lf)* mutants, the Ca^2+^ frequency was significantly increased to 5.9±0.5 Hz/300s. Yet, the Ca^2+^ amplitude of *twk-40(hp834)* was unchanged (**Figure 3G**), and the individual Ca^2+^ spike kinetics – including the rise and decay time – were not modified either (**Supplementary Figure S3A-C**), suggesting that *twk-40* contributes to the basal-activity of the DVB neuron. Taken together, our experiments demonstrate that the intrinsic DVB Ca^2+^ oscillation activity was substantially regulated by *twk-40*.

To further decipher *twk-40*’s cellular specificity, we firstly measured neuronal Ca^2+^ activity by expressing wild-type TWK-40 cDNA under different exogenous promoters in a *twk-40(bln336, gf)* mutant background. Consistent with our behavioral data, the Ca^2+^ frequency and amplitude were partially rescued when TWK-40(WT) was expressed in DVB in *twk-40(bln336, gf)* mutants (**Figure 4A-C**). By contrast, neither frequency nor amplitude was restored when TWK-40(WT) was expressed in D-motor neurons (**Figure 4C**). In addition, we then expressed TWK-40(WT) in *twk-40(hp834, lf)* mutant and found that the increased Ca^2+^ frequency of these mutants could also be rescued to wild-type levels when TWK-40(WT) was expressed in DVB (**Figure 4D-F**). Thus, TWK-40(WT) is sufficient to restore the neuronal Ca^2+^ activity of the DVB neurons.

**Figure 4.**
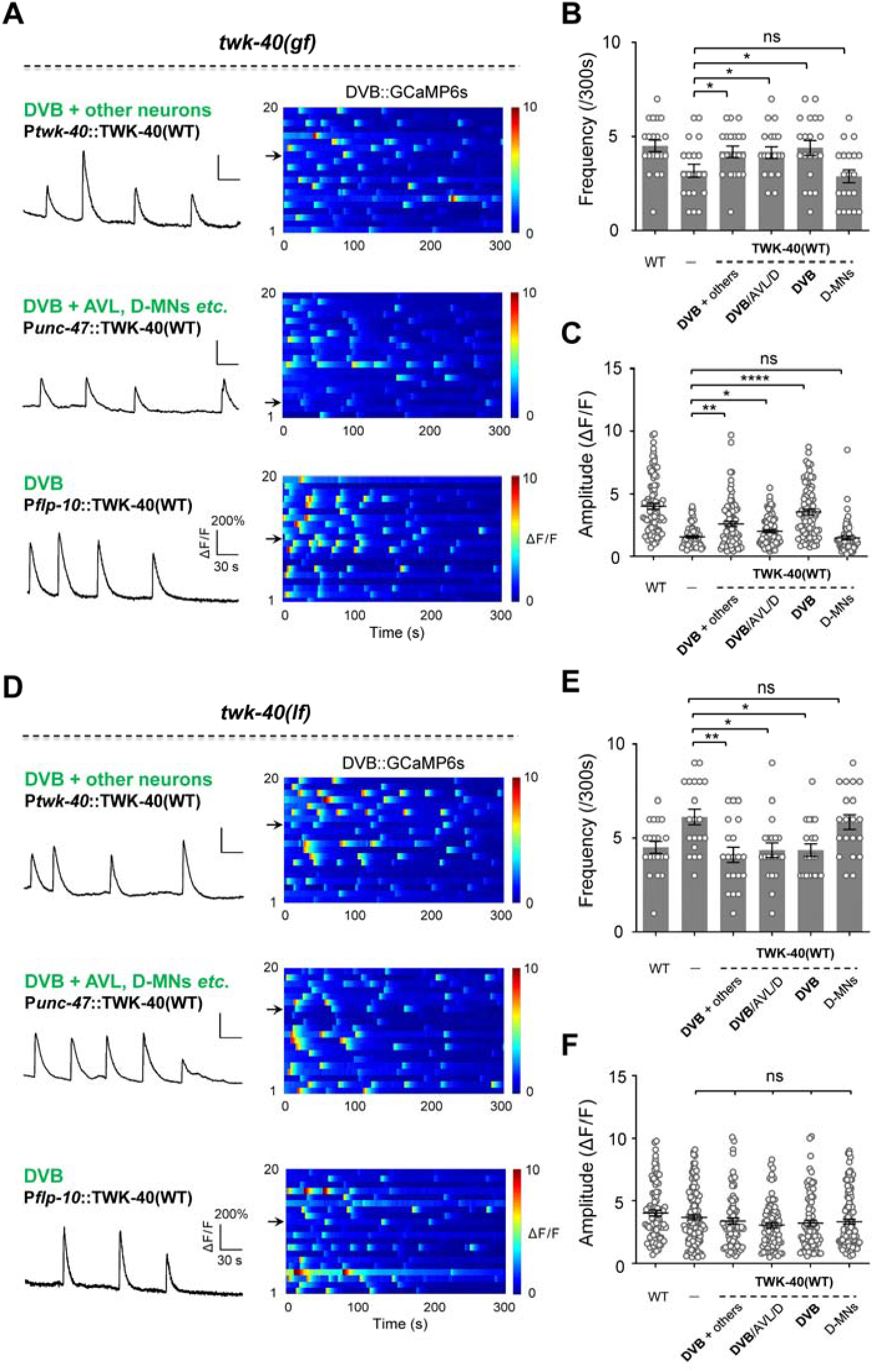
twk-40 regulates Ca^2+^ oscillation in DVB neuron. (**A**) Representative DVB Ca^2+^ traces (*left*) and color maps (*right*) for tissue-specific expression of TWK-40(WT) in *twk-40(bln336, gf)* mutants. *Upper*, endogenous promoter P*twk-40*; *middle*, P*unc-47* promoter including DVB expression; *bottom*, P*unc-25* promoter lacking DVB expression. (**B, C**) Quantification of the frequency and amplitude of Ca^2+^ transients, n=20. (**D**) Representative DVB Ca^2+^ traces (*left*) and color maps (*right*) for tissue-specific expression of TWK-40(WT) in *twk-40(hp834, lf)* mutant. (**E, F**) Quantification of the frequency and amplitude of Ca^2+^ transients, n=20 animals. All data are expressed as means ± SEM. One-way ANOVA test was used, in which: ns, not significant, \**P* < 0.05, \*\**P* < 0.01, \*\*\*\**P* < 0.0001 in comparison with that as denoted.

Collectively, these results support the notion that *twk-40* regulates neuronal Ca^2+^ oscillations of the DVB neuron.

### twk-40 cell-autonomously regulates DVB Ca^2+^ oscillations

The broad neuronal expression pattern of *twk-40* indicates that TWK-40 may regulate the DMP not limited in the DVB neuron (White *et al*., 1986). To examine whether synaptic input upstream of DVB may play a role, we examined the DVB Ca^2+^ oscillations in an *unc-13(lf)* background (Wang *et al*., 2013). UNC-13 is a conserved and essential presynaptic Ca^2+^ effector that triggers exocytosis and neurotransmitter release (Brose et al., 1995; Gao and Zhen, 2011; Richmond and Jorgensen, 1999). Loss of function of *unc-13* strongly impairs global neurotransmission with minor effect on NLP-40 release (Wang *et al*., 2013). Interestingly, in the *unc-13(lf)* mutant background, we found that Ca^2+^ oscillation defects were retained in *twk-40* mutants, including the decreased frequency and amplitude in *twk-40(bln336, gf)* mutants and increased frequency in *twk-40(hp834, lf)* mutants (**Supplementary Figure S4A-C**). These results demonstrate that potential presynaptic inputs to DVB don’t interfere with the regulation of DVB’s Ca^2+^ activities by TWK-40, and that *twk-40* most likely regulates the cellular excitability of DVB neuron in a cell-autonomous manner.

To reinforce this notion, we further tested TWK-40’s inhibition of DVB by measuring the Ca^2+^ oscillation in different transgenic lines that expressed the *twk-40(L159N, gf)* cDNA. When TWK-40(L159N) was expressed in DVB in wild-type animals (P*twk-40*, P*unc-47* or P*flp-10*), the frequency and amplitude of Ca^2+^ spikes were significantly reduced (**Supplementary Figure S5A**), similar to *twk-40(bln336, gf)* mutants (**Supplementary Figure S5B, C**). By contrast, expression of TWK-40(L159N) in D-motor neurons had no effect. Furthermore, reduced Ca^2+^ oscillations were also observed when TWK-40(L159N) was expressed in a *twk-40(hp834, lf)* mutant background (**Supplementary Figure S5E-F**). These results reveal that ectopic expression of TWK-40(L159N) is sufficient to silence the Ca^2+^ oscillation of DVB neurons. Namely, TWK-40 execute a dominant inhibition of the intrinsic activity regulation in DVB neuron.

### DVB activity regulates Ca^2+^ dynamics in enteric muscles

The expulsion behavior ultimately relies on the coordinated contraction of a group of muscles, including the anal depressor and sphincter, and two enteric muscles (EM), which wrap around the posterior gut to further pressurize the intestinal contents (McIntire *et al*., 1993b). EMs are innervated by DVB (**Figure 5A**) and AVL neurons (McIntire *et al*., 1993b; White *et al*., 1986), and the contraction of EMs is driven by intracellular Ca^2+^ transients (LeBoeuf and Garcia, 2017). We then ask whether EM Ca^2+^ activity is also regulated in *twk-40* mutants.

**Figure 5.**
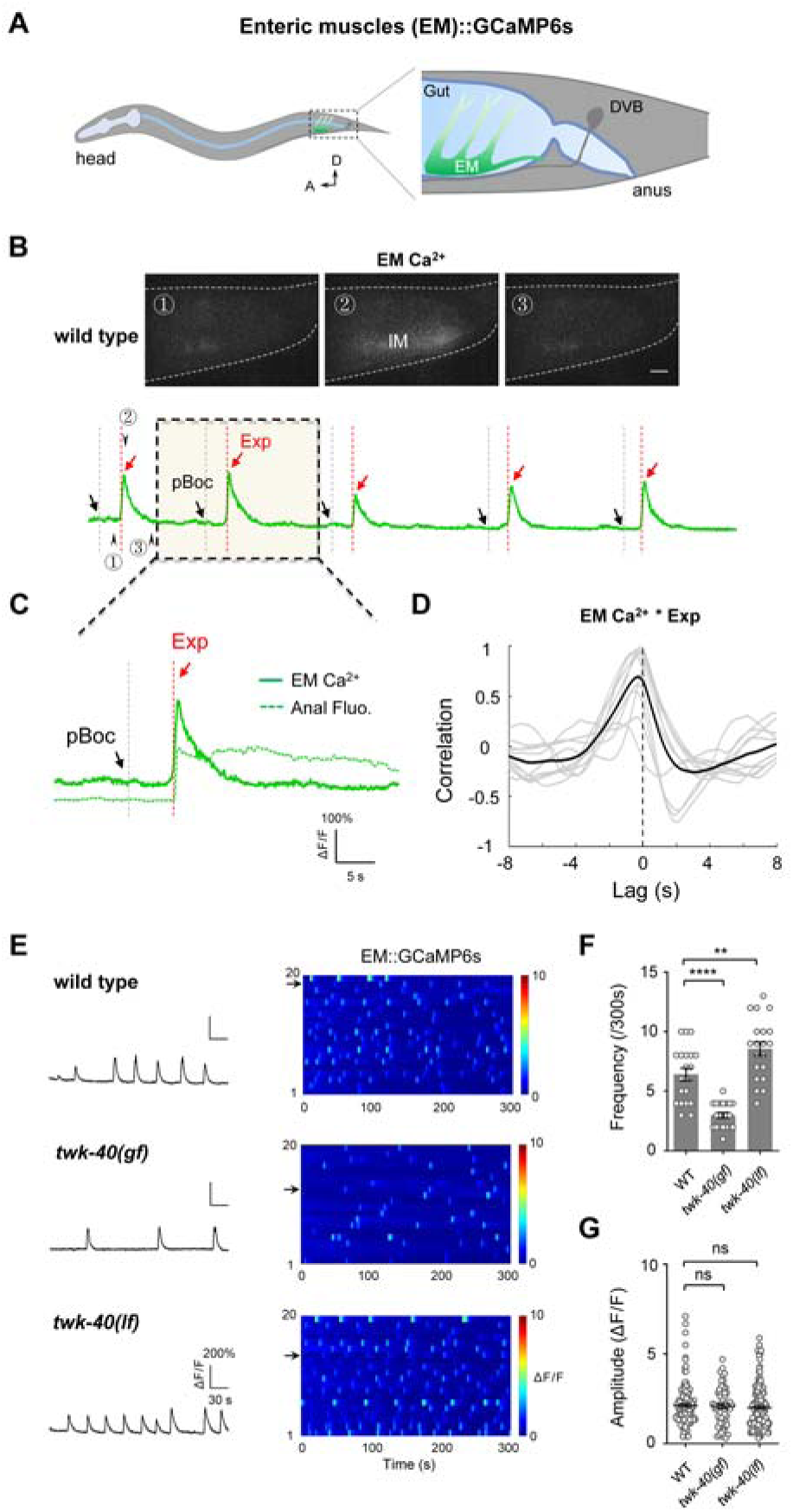
twk-40 mutants regulate Ca^2+^ dynamics of enteric muscles. (**A**) *i*. Schematic representation of *C. elegans* DVB neuron and enteric muscle (EM) in a lateral view. The genetically-encoded Ca^2+^ indicator GCaMP6s was expressed in EM. *ii*. Representative Ca^2+^ oscillation imaged by GCaMP in EM *in-vivo*. *Upper*, from left to right, Ca^2+^ signal of EM in (1) the quiescent state, (2) at peak Ca^2+^ signal and (3) upon return to the basal state. Scale bar, 5 μm. *Bottom*, representative recording of Ca^2+^ oscillations in EM over time. Arrow heads indicate the corresponding time points in upper panels. Color map of the rhythmic EM Ca^2+^ activities in wild-type animals. (**B**) Representative Ca^2+^ activity (*left*) and color maps (*right*) of enteric muscles in *twk-40(bln336, gf)* and *twk-40(hp834, lf)* mutants, respectively. (**C**) Quantification of EM Ca^2+^ transients frequency and amplitude, n=20. Ca^2+^ transient frequency was significantly reduced in *twk-40(bln336, gf)* mutant, but was increased in *twk-40(hp834, lf)* mutant. EM Ca^2+^ amplitude was not altered in either mutants. All data are expressed as means ± SEM. One-way ANOVA-test was used, in which: ns, not significant, \*\**P* < 0.05, \*\*\*\**P* < 0.0001 in comparison with that as denoted.

We thus performed Ca^2+^ imaging of animals expressing the calcium indicator GCaMP6s in the enteric muscles (EM::GCaMP6s, **Figure 5B**) (Li *et al*., 2023). Similar to activation dynamics in DVB neuron, enteric muscles displayed rhythmic Ca^2+^ spikes closely correlated with expulsion action (**Figure 5C, D**). In our experimental conditions, the EM Ca^2+^ spikes exhibited a frequency of 6.4±0.5 Hz/300s (n = 20) in wild-type animals (**Figure 5E**), which was also eliminated in *nlp-40(tm4085)* mutant (**Supplementary Figure S2D-E**). In *twk-40* mutants, the periodicity of EM Ca^2+^ spikes was affected. Specifically, the EM Ca^2+^ frequency was significantly reduced in *twk-40(bln336, gf)* mutants (3.0±0.2 Hz/300s, n = 20), and increased to 8.6±0.6 Hz/300s (n = 20) in *twk-40(hp834, lf)* mutants (**Figure 5E, F**), reminiscent of *twk-40*’s effects for DVB Ca^2+^ frequency. EM Ca^2+^ amplitudes, however, were unchanged in *twk-40(bln336, gf)* animals (**Figure 5G**), which differs from the reduction observed for DVB Ca^2+^ spike amplitudes. Given that no *twk-40* expression was observed in enteric muscles, we propose that the effects of *twk-40* mutations on the Ca^2+^ activity of enteric muscles are a consequence of the dysregulation of presynaptic DVB neurons.

To confirm this idea, we performed additional experiments by expressing wild-type TWK-40 in EM and DVB neurons, respectively. We found that the reduced EM Ca^2+^ frequency in *twk-40(bln336, gf)* mutant was almost fully rescued by expressing TWK-40(WT) in DVB (P*twk-40*, P*unc-47* and P*flp-10*), but not in enteric muscles (P*exp-1*) or in D-motor neurons (P*unc-25s*) (**Figure 6A, B**). Consistently, the increased frequency of EM Ca^2+^ spikes in *twk-40(hp834, lf)* mutants was also restored to wild-type by TWK-40(WT) expression in DVB neurons (**Figure 6C, D**). We also ectopically expressed the gain-of-function TWK-40(L159N) in *twk-40(hp834)* animals to inhibit DVB’s activity and recorded the EM Ca^2+^ spikes. Consistently, the frequency of EM Ca^2+^ spikes was significantly diminished in this experiment. This inhibition was however not observed in animals expressing TWK-40(L159N) in the enteric muscles or D-motor neurons (**Figure 6E, F**). Collectively, our experiments argue that *twk-40* regulates Ca^2+^ activities of enteric muscles by modulating DVB activity.

**Figure 6.**
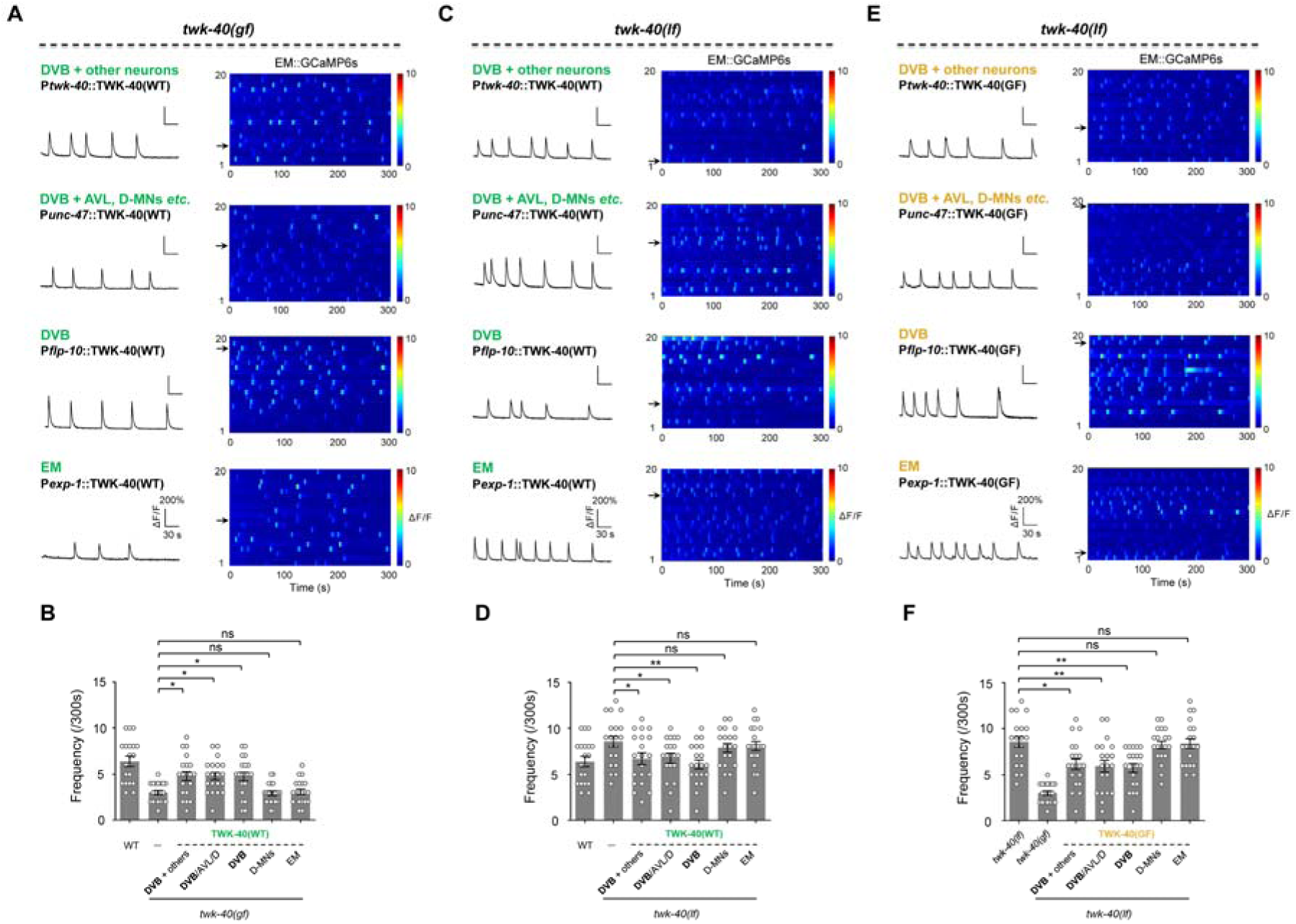
DVB regulates enteric muscles Ca^2+^ dynamics in twk-40 mutants. (**A, C**) Representative EM Ca^2+^ traces (*left*) and color maps (*right*) in *twk-40(bln336, gf)* (A) and *twk-40(hp834, lf)* (C) for tissue-specific expression of TWK-40(WT). 1, endogenous promoter P*twk-40*; 2, P*unc-47* promoter including DVB expression; 3, P*unc-25* promoter lacking DVB expression; 4, P*exp-1* promoter for enteric muscle expression. (**B, D**) Rescue of EM Ca^2+^ transient frequency by expression of TWK-40(WT) in DVB, but not enteric muscles or other neurons, n=20. (**E**) Representative EM Ca^2+^ traces (*left*) and color maps (*right*) in *twk-40(hp834, lf)* following tissue-specific expression of TWK-40(L159N). (**F**) Reduction of EM Ca^2+^ transient frequency following TWK-40(L159N) expression in DVB, but not in enteric muscles or other neurons, n=20. All data are expressed as means ± SEM. One-way ANOVA test was used, in which: ns, not significant, \**P* < 0.05, \*\**P* < 0.01in comparison with that as denoted.

### TWK-40 hyperpolarizes the resting membrane potential of DVB neuron

The functional inhibition of DVB neuron by *twk-40* is consistent with an inhibitory function of this potassium channel. To directly investigate the functional effect of *twk-40*, we dissected and recorded GFP-labeled DVB neurons *in vivo* in the whole-cell patch clamp configuration (Methods, **Figure 7A**). In wild-type animals, stable outward K^+^ currents were recorded at stepwise holding voltages from -60 mV to +80 mV (Jiang et al., 2022; Xie et al., 2013). These currents were significantly diminished in *twk-40(hp834)* loss-of-function mutants (**Figure 7B, C**). Conversely, in *twk-40(bln336)* gain-of-function mutants, outward currents were dramatically increased (**Figure 7B, C**). Thus, these *in-situ* neuronal recordings demonstrate that *twk-40* supports a functional K^+^ current in DVB neuron.

**Figure 7.**
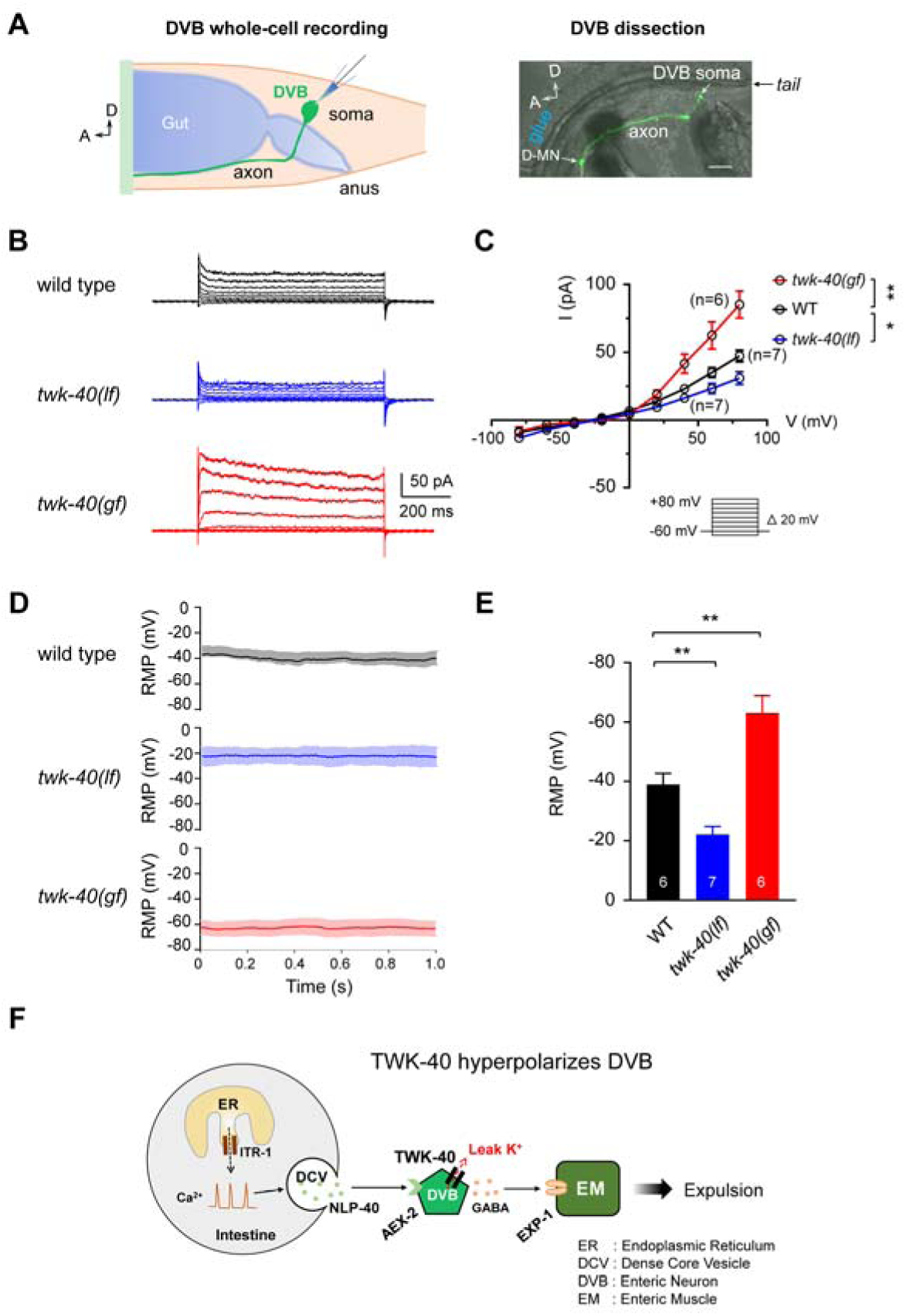
TWK-40 hyperpolarizes the DVB neuron. (**A**) *Left*, schematic representation of *C. elegans* DVB neuron for whole-cell patch clamp recording. *Right*, GFP-labeled DVB soma and axon. (**B**) Representative outward K^+^ currents recorded from DVB neurons in different genotypes. (**C**) Quantification of the current-voltage curves from different genotypes reveals that the outward currents were reduced in *twk-40(hp834, lf)* mutant, but increased in *twk-40(bln336, gf)* mutant. (**D, E**) Representative traces and quantification of the resting membrane potential (RMP) recorded from DVB in different genotypes. (**F**) The working Model: K_2P_ channel TWK-40 regulates the rhythmic expulsion behavior by setting the resting membrane potential of DVB neuron. One-way ANOVA test was used to test the significant difference of average currents at +80 mV, in which: ns, not significant, \**P* < 0.05, \*\**P* < 0.01 in comparison with that as denoted.

Furthermore, the resting membrane potential of DVB was also analyzed. While wild-type DVB neurons had an RMP of -38.9±3.7 mV (n = 6), loss of *twk-40* depolarized the RMP to -22.1±2.9 mV (n = 7). Conversely, in *twk-40(bln336)* mutants, the RMP of DVB neuron was dramatically hyperpolarized to -63.1±5.8 mV (n = 6) (**Figure 7D, E**).

Taken together, these results demonstrate that TWK-40 constitutes a K^+^ channel that stabilizes the RMP of DVB neuron.

In summary, we show here that TWK-40 forms a *bona fide* potassium channel that cell-autonomously maintains the RMP of DVB neuron, and that *twk-40* is essential for the regulation of the rhythmic expulsion motor program (**Figure 7F**).

## Discussion

Potassium currents mediated by K_2P_ channels are important modulators of neuronal activity in animal nervous systems. We show here that the previously uncharacterized K_2P_ channel TWK-40 provides a prominent leak potassium current in the DVB motor neuron that regulates the rhythmic expulsion behaviour of *C. elegans*. We present genetic evidence that loss of *twk-40* suppresses the *exp* defects of *nlp-40* and *aex-2* mutants.

Additional evidence from expression pattern analysis, behavioral rescue experiments, and real-time neuronal and muscular Ca^2+^ imaging, reveal the substantial contribution of TWK-40 to rhythmic expulsion activity. Further proof collected by heterologous expression of TWK-40 (Meng *et al*., 2022) and *in situ* whole-cell patch clamp recording, demonstrates that *twk-40* constitutes a K_2P_-like potassium-selective current. Our results thus identify a K_2P_ channel TWK-40, through its influence on DVB neuron resting membrane potential, plays a critical role in coordinating the expulsion rhythm and locomotion, shedding light on the intricate interplay between potassium channels and neural circuits in controlling complex motor programs in vivo. These findings suggest a broader significance of TWK-40/K_2P_ in orchestrating multiple motor rhythms, providing insights into the molecular mechanisms underlying coordinated motor behaviors.

Circadian rhythms with 24-hour-long cycles like sleeping are well studied and are controlled by the expression fluctuation of clock genes, like *Per1*, *Per2*,and *Bmal1 etc*. (Lowrey and Takahashi, 2004). Ultradian rhythms (or episodic ultradian events) with diverse shorter periods ranging from milliseconds to hours, are ubiquitous and indispensable in many neuromuscular systems, such as heartbeat, respiration and peristalsis (Goh et al., 2019). These ultradian rhythms are generally considered to be generated by motor central pattern generators (CPG) (Marder and Bucher, 2001). The ultradian DMP rhythm in *C. elegans* however is a remarkable exception, as it is initiated by calcium oscillations in intestinal cells (Teramoto and Iwasaki, 2006). The DVB and AVL neurons, acting as a signaling bridge through NLP-40 and GABA pathways, transfer non-neuronal calcium oscillation from the intestine to enteric muscles to execute the expulsion action.

Genetic analysis initially highlighted critical ion channel genes for DVB activity, such as *unc-2* and *egl-19* that encode the α1 subunits of P/Q-and L-type voltage-gated Ca^2+^ channel, respectively (Wang and Sieburth, 2013), and *egl-36*, a Shaw-type (Kv3) voltage-dependent potassium channel subunit (Johnstone et al., 1997). However, the *in vivo* cellular mechanism of how these channels contribute to the behavior is still unknown. The recent descriptions of synchronized giant action potentials between AVL and DVB, and of unusual compound action potentials in AVL, provide insights into how the rhythmic expulsion behaviour is generated at the cellular level. In particular, the negative potassium spikes in AVL are mediated by a repolarization-activated potassium channel EXP-2 (Jiang *et al*., 2022). Yet, no ion channel had been associated so far with the rhythmic activity of DVB. Thus, this study identifies TWK-40 as a novel important electrical component for the regulation of the defecation motor program.

Among 47 K_2P_ genes in the *C. elegans* genome (Bargmann, 1998), only few have been substantially investigated. Interestingly, nematode K_2P_ channels have been mostly studied using gain-of-function mutations identified in forward genetic screens. For instance, while *twk-18(gf)* mutants cause sluggish, uncoordinated movement (Kunkel et al., 2000), *sup-9(gf)* mutants are hyperactive with a characteristic rubberband phenotype (de la Cruz et al., 2014). Gain-of-function mutants of *unc-58* show significant deficits in locomotion, egg laying, development and aging (Kasap et al., 2018; Salkoff et al., 2005). Except for *twk-18*, we still lack a comprehensive understanding of the electrical properties of these channels *in vitro* and *in vivo*. The recent discovery of a universal activating mutation that promotes the gating of vertebrate and invertebrate K_2P_ channels has opened the way to more comprehensive manipulation of ion channel activity (Ben Soussia *et al*., 2019). We used this strategy here to engineer a point mutation of TWK-40 L159N that achieved a substantial gain-of-function effect. First, we show in a related study that the currents conducted by TWK-40(L159N) channels exhibit ∼5 fold increase compared to TWK-40(WT) when heterologously expressed in HEK293 cells (Meng *et al*., 2022). Second, the resting membrane potential of DVB neurons was drastically hyperpolarized due to the increased outward K^+^ current. In this *twk-40(bln336)* gain-of-function mutant, the neuronal Ca^2+^ oscillation frequency and amplitude were reduced, resulting in the significantly diminished expulsion frequency. Thus, our study not only identifies TWK-40 as a regulator of the expulsion rhythm and DVB RMP, but also provides insight into the electrical properties of this K_2P_ channel.

We also describe the impact of loss of *twk-40* on the expulsion behavior and DVB activity. In contrast to *gf* mutants, the resting membrane potential of DVB neuron was depolarized in *twk-40(lf)* mutant, and high Ca^2+^ oscillation activity was observed. Consequently, the expulsion number exhibited a moderate but significant increase in *twk-40(lf)* mutant. The fact that only few K_2P_ mutants with loss of function phenotypes have been identified in *C. elegans* may be due to functional redundancy between K_2P_ genes. *twk-7* is a notable exception as it exhibits hyperactive locomotion (Gottschling et al., 2017; Luersen et al., 2016). Interestingly, we have found that *twk-40(lf)* mutants also exhibit increased forward and backward body bends, as well as more frequent and prolonged reversals (Meng *et al*., 2022). A recent genetic study reported that loss of function *twk-40* could reverse the reduced body curvature of *C. elegans NALCN(lf)* mutants (Zhou et al., 2022). Thus, TWK-40 appears to be required for the regulation of multiple motor rhythms.

The slight outward-rectification of the *twk-40* dependent K^+^ currents in DVB neurons differs from the voltage-independent currents observed in recombinant TWK-40 recorded in HEK293 cells (Meng *et al*., 2022). This could be due to endogenous regulatory subunits expressed in DVB neurons that modify the channel kinetics of TWK-40. Alternatively, TWK-40 could form heteromers with other K_2P_ subunits *in vivo*, resulting in heterodimers with different activation kinetics (Czirjak and Enyedi, 2002). K_2P_ channels can be modulated by a variety of biophysical parameters, such as pH, temperature, and mechanic forces (Enyedi and Czirjak, 2010). Certain K_2P_ channels, including TASK-1/3, TREK-1, and TRESK, are activated by volatile general anesthetics at clinically relevant concentrations (Liu et al., 2004; Patel et al., 1999; Sirois et al., 2000), indicating a major class of drug target from these K_2P_ channels. As a possible ortholog of human TASK-1/3, identification of the critical stimulator of TWK-40 in the future could promote the understanding of how the external environment and internal state affect motor rhythm.

## Materials and Methods

### C. elegans strains and transgenic lines

The complete lists of constructs, transgenic lines and strains generated or acquired for this study are provided in the Supplementary Table 1 and Table 2. All *C. elegans* strains were cultured on standard Nematode Growth Medium (NGM) plates seeded with OP50, and maintained at 22 °C. Multiple extrachromosomal transgenic lines were generated for all rescue experiments and visually examined for consistency. One representative transgenic line was quantified and results shown in most experiments. Unless stated otherwise noted, the wild-type animal refers to the Bristol N2 strain. Only hermaphrodite worms were used for the experiments.

### Constructs and molecular biology

All expression plasmids in this study were constructed by Three-Fragment Multisite Gateway (Invitrogen, Thermo Fisher Scientific, USA) (Magnani et al., 2006). Three-Fragment Multisite gateway system consists of three entry clones. Three entry clones comprising three PCR products (promoter, target gene, sl2-GFP or unc-54 3’UTR, in name of slot 1, slot 2 and slot 3, respectively) were recombined into the pDEST-R4-R3 or pDEST-R4-R3-unc-54 3’UTR destination vectors by using standard LR recombination reactions to generate the expression clones.

All promoters used in this study were generated by PCR against a mixed-stage N2 *C. elegans* genomic DNA. Promoters include the 1.4 kb P*twk-40*, 1.2 kb P*unc-47*, 1.6 kb P*flp-10*, 1.8 kb P*unc-25s*, 2.4 kb P*exp-1* and 1.4 kb P*ges-1* genomic sequence upstream of the respective ATG start codon and were used to substitute for the *rab-3* fragment in standard BP reaction-generated entry clone A with the In-Fusion method, using ClonExpress®ⅡOne Step Cloning Kit (Vazyme, China).

To generate the entry clones slot 2 and slot 3, we used standard BP recombination reactions. An entry clone B contributing sequences of slot 2 in the expression plasmid contains sequence of a target gene, i.e., *twk-40*, GFP, or RFP. The “B” entry clones were constructed by the use of *pDONR™ 221* donor vector (Invitrogen) through BP reactions. All the DNA fragments used to construct entry clone B involved in this project were amplified by PCR with the primers containing attB1 and attB2 recombination sites. An entry clone “C” contributing sequences of slot 3 in the expression plasmids contains a sequence of *unc-54* 3’ UTR, *sl2d*-GFP or *sl2d*-wCherry. The “C” entry clones were constructed by use of *pDONR-P2R-P3* donor vector through standard BP reactions. The corresponding PCR products with attB2R and attB3 sites were amplified by PCR.

For calcium imaging, constructs with genetic calcium sensor GCaMP6s were used for neuronal activity measurement. The GCaMP6s sequence (Chen et al., 2013) was codon-optimized for expression in *C. elegans*. The synthesized gene also contains three *C. elegans* introns and contain restriction enzyme sites to facilitate subsequent cloning. In all constructs, GCaMP6s was fused in frame with a *C. elegans* codon-optimized wCherry at the C-terminus.

### Genetic mutants and transgenetic arrays

Loss-of-function allele *hp834* was generated by targeted CRISPR/Cas9 genome editing with a 7 bp deletion of the sequence (GTTCGAG, base 127-133) was introduced in the 2nd exon of T28A8.1a. *twk-40(bln336)* was generated by CRISPR/Cas9 technology by mutating 1047-1048 base of T28A8.1a from CT to AA. Other genetic mutants used for constructing transgenic lines and compound mutants were obtained from the *Caenorhabditis Genetics Center* (*CGC*). Transgenic animals that carry non-integrated, extra-chromosomal arrays (*gaaEx*) were generated by co-injecting an injection marker with one to multiple DNA construct at 5-30 ng/μl. All animals were backcrossed at least four times against N2 prior to analyses.

### Behavioral assay for the defecation motor program

Defecation behavior was measured on NGM plates under standard conditions as previously described (Thomas, 1990). All behavioral tests were performed at 20-22°C. Worms were synchronized and well-fed young hermaphrodites adults were transferred to fresh NGM plates with OP50 for behavior assay during freely-moving state. The defecation phenotypes were scored under a stereo fluorescence microscope (Zeiss V16, Germany). Each animal was quantified in 10 minutes after the first pBoc step pass using the Etho program software (Liu and Thomas, 1994). We simplified the observation on the pBoc and the Exp steps, due to the ambiguous aBoc step (Iwasaki and Thomas, 1997). Exp/cycle was calculated as the ratio of the number of Exp/pBoc during a 10 min recording (Wang *et al*., 2013).

### Fluorescence Microscopy

Images of fluorescently tagged proteins were captured in live L4 worms. L4 stage transgenic animals expressing fluorescence markers were picked a day before imaging. Worms were immobilized by 2.5 mM levamisole (Sigma-Aldrich, USA) in M9 buffer. Fluorescence signals were captured from live worms using a Plan-Apochromatic 60X objective on a confocal microscope (FV3000, Olympus, Japan) in the same conditions.

### Calcium imaging

The two integrated strains *hpIs468* (P*unc-47*::GCaMP6s::wCherry) and *hpIs582* (P*exp-1*::GCaMP6s::wCherry) were used for calcium imaging of DVB neuron and enteric muscles, respectively. Under standard OP50 feeding, two strains exhibit normal defecation interval and Exp step. To simplify the experiment, young hermaphrodite adult animals were glued dorsally as described previously (Gao et al., 2015), and imaged with a 60x water objective (Nikon, Japan, numerical aperture = 1.0) (Li *et al*., 2023). In particular, animals are loosely immobilized with WormGlu (Histoacryl Blue, Braun, Germany) to a sylgard-coated cover glass covered with bath-solution (Sylgard 184, Dowcorning, USA) but leave the tail free in the recording solution. To keep the worms’ regular pumping and defecation activities, a small amount of OP50 (∼100 µl, OD600 = 1) was mixed into the recording solution, which consists of (in mM): NaCl 150; KCl 5; CaCl_2_ 5; MgCl_2_ 1; glucose 10; sucrose 5; HEPES 15, pH7.3 with NaOH, ∼330 mOsm.

Fluorescence images were acquired with excitation wavelength LED at 470 nm (LED, BioLED Light Source Control Module 9-24V DC) and a digital sCMOS camera (Hamamatsu ORCA-Flash 4.0 V2) at 100 ms per frame for 3 minute. Data was collected from HCImage (Hamamatsu) and analyzed by Image Pro Plus 6.0 (Media Cybernetics, Inc., Rockville, MD, USA) and Image J (National Institutes of Health). The GCaMP6s fluorescence intensity of the region of interest (ROI) was defined as F, the background intensity near the ROI was defined as F_0_. The true neuronal calcium fluorescence signal was obtained by subtracting the background signal from the ROI. ΔF/F_0_ = (F-F_0_)/F_0_ was plotted over time as a fluorescence variation curve.

### In situ electrophysiology

Dissection and recording were carried out using protocols and solutions described in the previous studies (Gao and Zhen, 2011; Richmond and Jorgensen, 1999). Briefly, 1-or 2-day-old hermaphrodite adults were glued (Histoacryl Blue, Braun, Germany) to a sylgard-coated cover glass covered with bath solution (Sylgard 184, Dowcorning, USA) under stereoscopic microscope M50 (Leica, Germany). After clearing the viscera by suction through a glass pipette, the tail cuticle flap was turned and gently glued down using WORMGLU (GluStitch Inc., Canada) to expose the DVB soma from *gaaEx0112* [P*unc-47*::GFP::3’UTR; P*myo-2*::GFP]. The integrity of the ventral nerve cord and DVB neuron were visually examined via DIC microscopy (Eclipse FN1, Nikon, Japan), and DVB soma were patched using 15-20 MΩ-resistant borosilicate pipettes (1B100F-4, World Precision Instruments, USA). Pipettes were pulled by micropipette puller P-1000 (Sutter, USA), and fire-polished by microforge MF-830 (Narishige, Japan). Membrane potentials and currents were recorded in the whole-cell configuration by EPC9 amplifier (HEKA, Germany), and analyzed using the Igor Pro (WaveMetrics, USA) and Clampfit 10 software (Axon Instruments, Molecular Devices, USA). Membrane potential was recorded at 0 pA, I-V curve was collected at holding potential from –60 mV to +80 mV at 20 mV increment. Data were digitized at 10-20 kHz and filtered at 2.6 kHz. The pipette solution contains (in mM): K-gluconate 115; KCl 25; CaCl_2_ 0.1; MgCl_2_ 5; BAPTA 1; HEPES 10; Na_2_ATP 5; Na_2_GTP 0.5; cAMP 0.5; cGMP 0.5, pH7.2 with KOH, ∼320 mOsm. cAMP and cGMP were included to maintain the activity and longevity of the preparation. The bath solution consists of (in mM): NaCl 150; KCl 5; CaCl_2_ 5; MgCl_2_ 1; glucose 10; sucrose 5; HEPES 15, pH7.3 with NaOH, ∼330 mOsm. Chemicals were obtained from Sigma unless stated otherwise. Experiments were performed at room temperatures (20-22 °C).

### Statistical analysis

The two-tailed Student’s *t*-test were used to compare two groups data sets. When multiple groups (more than two groups) of data were compared, ordinary one-way or two-way ANOVA analysis was performed for statistical analysis. The Shapiro-Wilk test was used to check for normality and either Levene’s test to check for equal variance. *P* < 0.05 was considered to be statistically significant; *, **, *** and **** denote *P* < 0.05, *P* < 0.01, *P* < 0.001, *P* < 0.0001, respectively. Graphing and subsequent analysis were performed using Igor Pro (WaveMetrics), Clampfit (Molecular Devices), Image J (National Institutes of Health), Python, Matlab (MathWorks, USA), GraphPad Prism 8 (GraphPad Software Inc., USA) and Excel (Microsoft, USA). For behavior analysis, calcium imaging, electrophysiology and fluorescence imaging, each recording trace was obtained from a different animal. Unless specified otherwise, data were presented as the mean ± SEM.

## Author Contributions

S.G. conceived experiments and wrote the manuscript. Z.Y., B.Y., Y.L., and Y.X. performed experiments and analyzed data. L.C., J.C. J.M., W.H., Y.T., S.E.M., and C.Z. contributed to the experiments. T.B. and M.Z. provided reagents, discussed the experiments and edited the text.

## Acknowledgements

We thank Chenhong Li, Long Ma for discussion. This research was supported by the Major International (Regional) Joint Research Project (32020103007), the National Natural Science Foundation of China (32371189, 31871069), the National Key Research and Development Program of China (2022YFA1206001), the Overseas High-level Talents Introduction Program, and the European Research Council (TB, ERC Starting Grant, Kelegans). We thank *Caenorhabditis Genetics Center*, which is funded by the NIH Office of Research Infrastructure Programs (P40 OD010440), for strains.

---

**Supplementary Figure 1.**
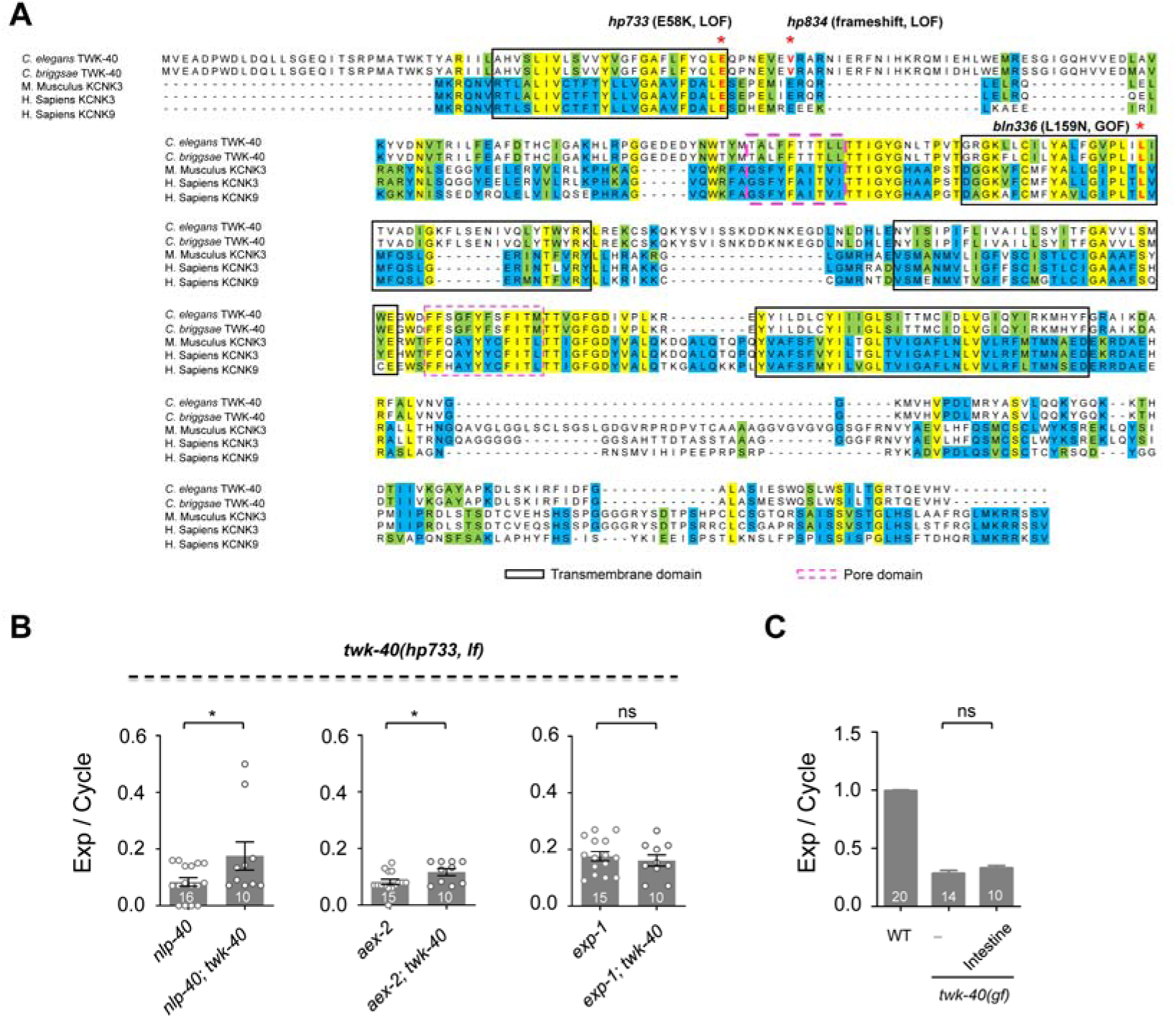
TWK-40 conservation, domain structure and gene mutations. (**A**) TWK-40 amino acid sequence alignment with other species. Identical amino acids are highlighted in yellow. Black boxes denote the transmembrane domains, pink dashed boxes denote the pore domains. Gene mutations are labeled on top of the sequence. (**B**) TWK-40 E58K suppresses the expulsion frequency per defecation cycle defects of *nlp-40* and *aex-2* mutants, but not *exp-1* mutants. (**C**) Expression of TWK-40(WT) in the intestine does not rescue the expulsion defect of *twk-40(bln336, gf)*. All data are expressed as means ± SEM. Student’s *t*-test was used, in which: ns, not significant, \**P* < 0.05, in comparison with that as denoted.

**Supplementary Figure 2.**
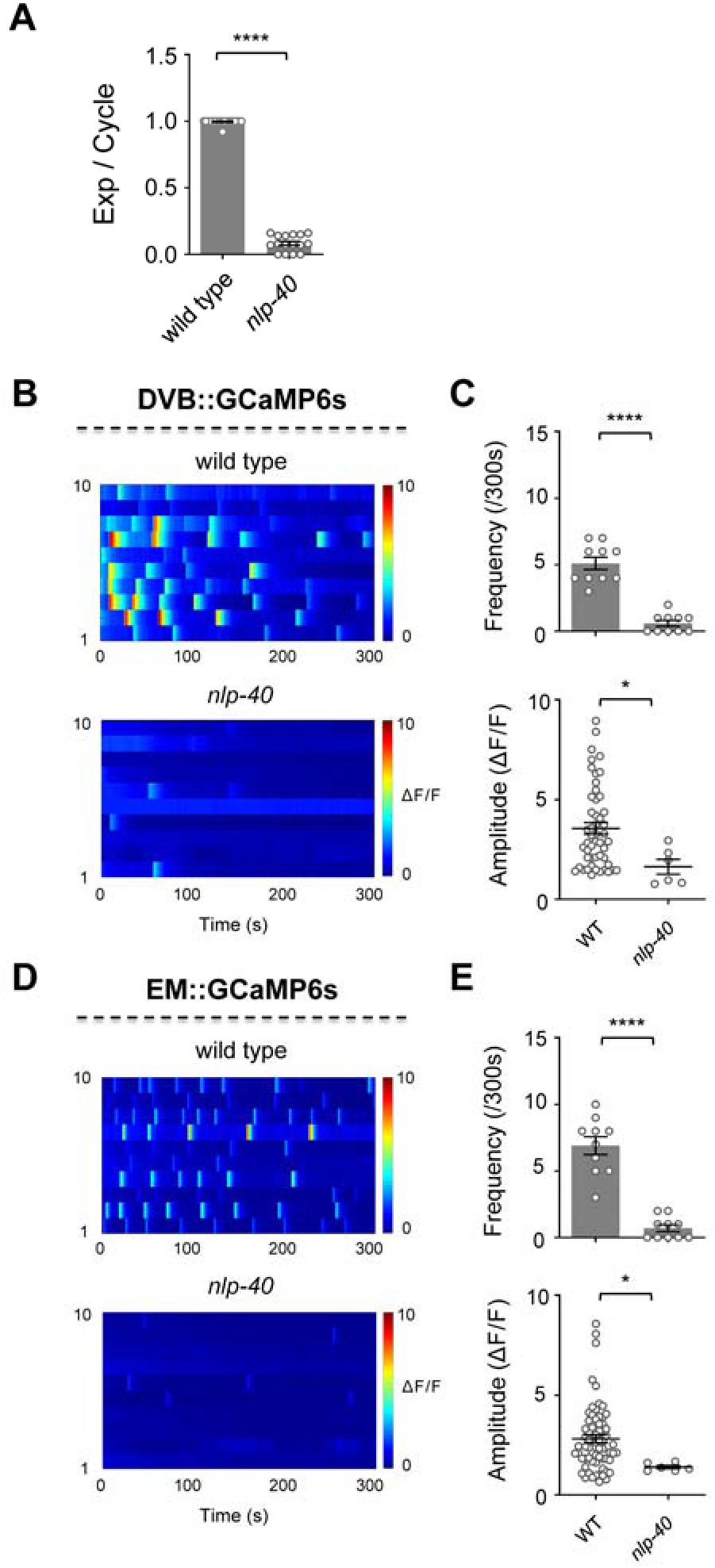
DVB/EM Ca^2+^ activities are dependent upon nlp-40. (**A**) Quantification of the expulsion frequency in *nlp-40(tm4085)* mutant. (B, D) Color maps summarize the DVB Ca^2+^ and EM Ca^2+^ activities in wild type and *nlp-40* animals, respectively. (C, E) Quantification of the frequency and peak amplitude of these Ca^2+^ activities. \**P* < 0.05; \*\*\*\**P* < 0.0001; two-tailed unpaired Student’s t-test. n = 10-30 animals. Error bars, SEM.

**Supplementary Figure 3.**
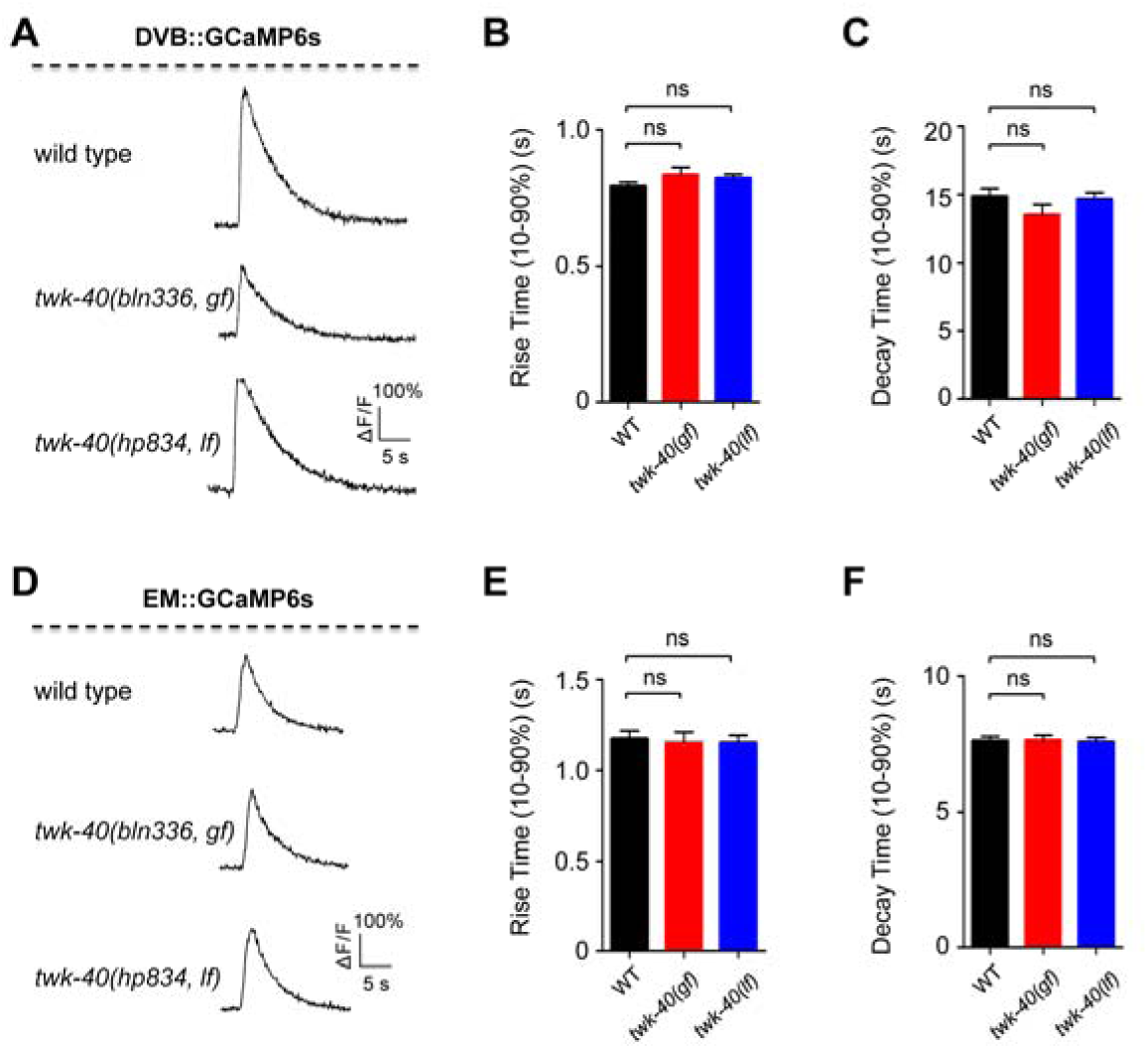
twk-40 *mutation does not* alter Ca^2+^ dynamics in DVB and enteric muscles. (**A, D**) Representative single Ca^2+^ transients from DVB (A) and enteric muscle (D) in wild type, *twk-40(bln336, gf)* and *twk-40(hp834, lf)* animals, respectively. (**B, C, E, F**) The rise time (10-90%) and decay time (10-90%) of the Ca^2+^ transients exhibited no difference between these genotypes. Each group, n=20 animals. All data are expressed as means ± SEM. One-way ANOVA test was used, in which: ns, not significant, in comparison with that as denoted.

**Supplementary Figure 4.**
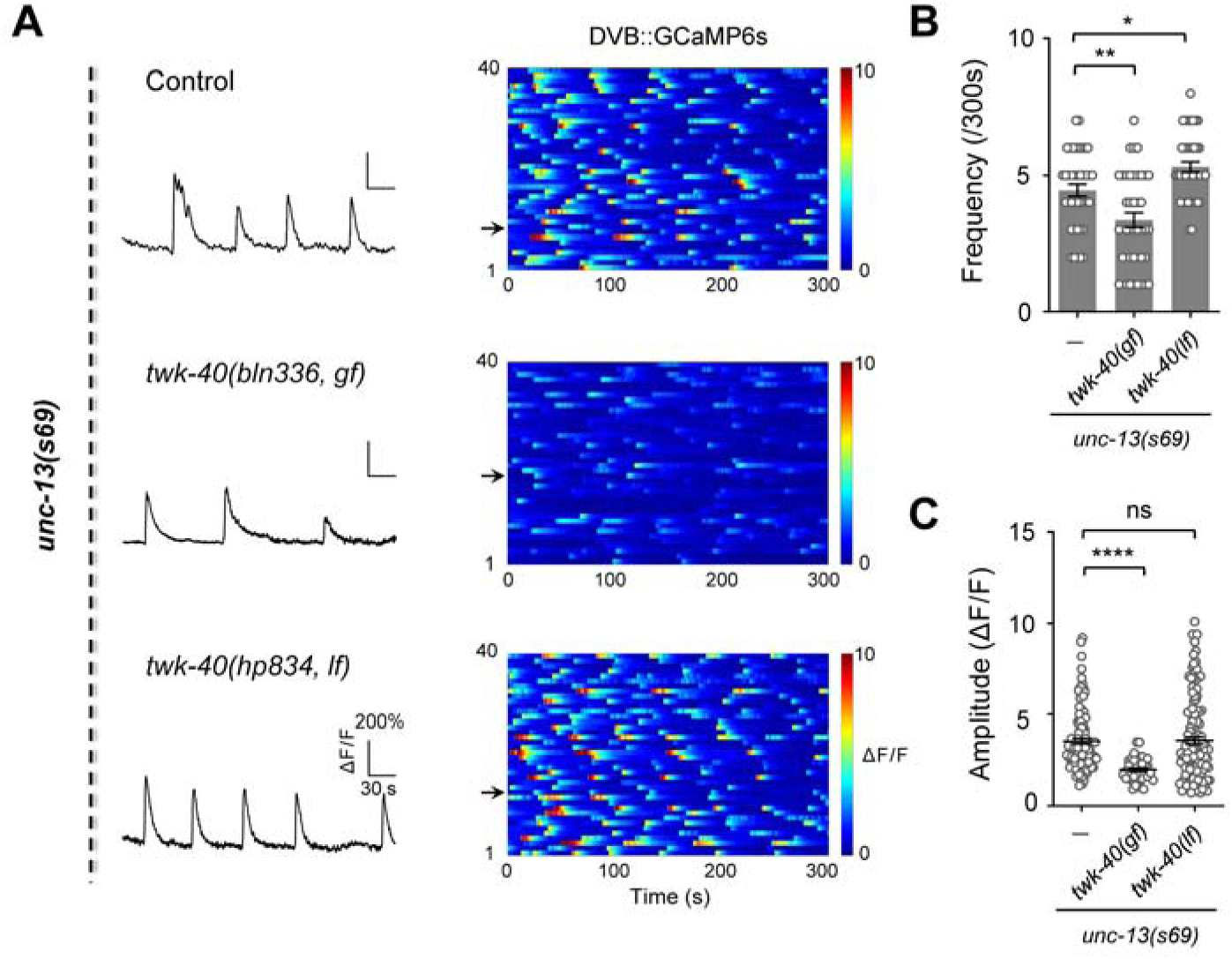
twk-40 regulates DVB Ca^2+^ oscillations in unc-13(lf) background. (**A**) Representative DVB recordings (*left*) and color maps of rhythmic Ca^2+^ oscillations (*right*) in *unc-13(s69)*, *unc-13(s69); twk-40(bln336, gf),* and *unc-13(s69); twk-40(hp834, lf)* mutants, respectively. (**B, C**) Quantification of the frequency and amplitude of DVB Ca^2+^ transients in different mutants. The frequency and amplitude were significantly reduced in *unc-13(s69); twk-40(bln336, gf)* mutants compared to *unc-13(s69)*. The Ca^2+^ frequency was slightly increased in *twk-40(hp834, lf)* mutant, but the amplitude was not altered. Each group, n=40. All data are expressed as means ± SEM. One-way ANOVA test was used, in which: ns, not significant, \**P* < 0.05, \*\**P* < 0.01, \*\*\*\**P* < 0.0001 in comparison with that as denoted.

**Supplementary Figure 5.**
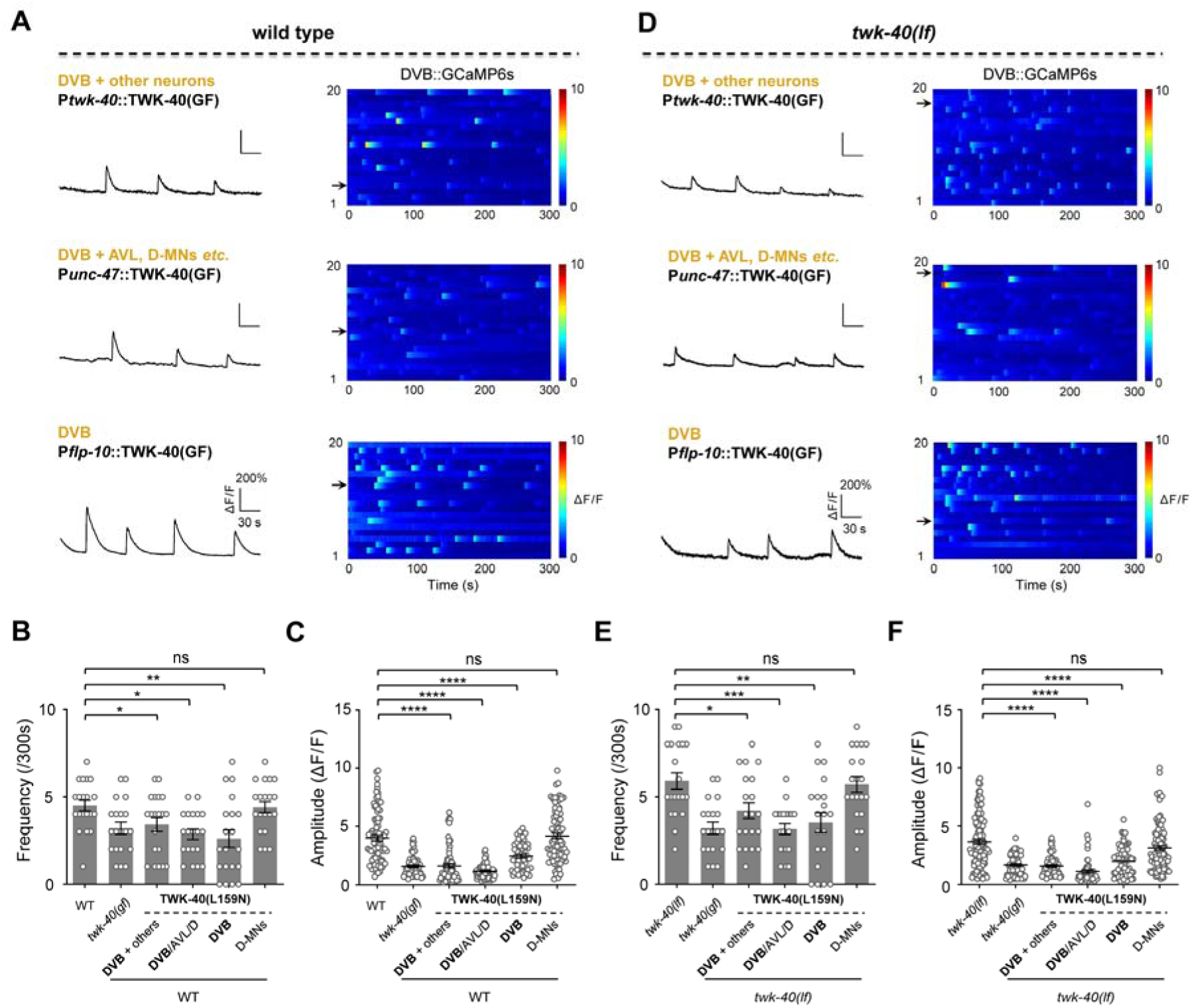
Ectopic expression of TWK-40(L159N) inhibits DVB. (**A, D**) Representative Ca^2+^ traces (*left*) and color maps (*right*) for DVB neurons in wild type (A) and *twk-40(hp834, lf)* (D) following expression of TWK-40(L159N) in different neuron groups. *Upper*, endogenous *twk-40* promoter; *middle*, *unc-47* promoter including DVB expression; *bottom*, short *unc-25* promoter (P*unc-25s*) lacking DVB expression. (**B, C, E, F**) Quantification of the frequency and amplitude of the oscillated DVB Ca^2+^ transients in different genetic backgrounds, both of which were reduced when TWK-40(L159N) was expressed in the DVB, but not in other neurons, n=20. All data are expressed as means ± SEM. One-way ANOVA test was used, in which: ns, not significant, \**P* < 0.05, \*\**P* < 0.01, \*\*\**P* < 0.001, \*\*\*\**P* < 0.0001 in comparison with that as denoted.

**Supplementary Table 1.**

| Strains | Genotype | Source |
| --- | --- | --- |
| ZM9786 | *twk-40(bln336, gf)* | M. Zhen Lab |
| ZM8302 | *twk-40(hp834, lf)* | M. Zhen Lab |
| JT3 | *aex-2(sa3)* | CGC |
| NM4394 | *nlp-40(tm4085)* | CGC |
| JT6 | *exp-1(sa6)* | CGC |
| ZM8661 | *unc-13(s69)* | M. Zhen Lab |
| ZM8518 | *hpIs468 [Punc-47::GCaMP6s:wCherry]* | M. Zhen Lab |
| ZM9095 | *hpIs582 [Pexp-1::GCaMP6s::wCherry]* | M. Zhen Lab |
| SGA359 | *aex-2(sa3); twk-40(hp834)* | This study |
| SGA360 | *nlp-40(tm4085); twk-40(hp834)* | This study |
| SGA372 | *exp-1(sa6); twk-40(hp834)* | This study |
| SGA370 | *aex-2(sa3); twk-40(bln336)* | This study |
| SGA371 | *nlp-40(tm4085); twk-40(bln336)* | This study |
| SGA373 | *exp-1(sa6); twk-40(bln336)* | This study |
| SGA374 | *aex-2(sa3); hpIs468* | This study |
| SGA375 | *aex-2(sa3); twk-40(hp834); hpIs468* | This study |
| SGA277 | *aex-2(sa3); twk-40(bln336); hpIs468* | This study |
| SGA331 | *unc-13(s69); hpIs468* | This study |
| SGA332 | *unc-13(s69); twk-40(hp834); hpIs468* | This study |
| SGA376 | *unc-13(s69); twk-40(bln336); hpIs468* | This study |
| SGA85 | *twk-40(bln336); hpIs468* | This study |
| SGA84 | *twk-40(bln336, gf); hpIs582* | This study |
| SGA83 | *twk-40(hp834); hpIs468* | This study |
| SGA82 | *twk-40(hp834); hpIs582* | This study |
| SGA61 | *twk-40(bln336); gaaEx061 [Ptwk-40::TWK-40(WT)::sl2dGFP]* | This study |
| SGA62 | *twk-40(bln336); gaaEx062 [Punc-47::TWK-40(WT)::sl2dGFP]* | This study |
| SGA71 | *twk-40(bln336); gaaEx071 [Punc-25::TWK-40(WT)::sl2dGFP]* | This study |
| SGA63 | *N<sub>2</sub>; gaaEx063 [Ptwk-40::TWK-40(GF)::sl2dGFP]* | This study |
| SGA78 | *N<sub>2</sub>; gaaEx078 [Punc-47::TWK-40(GF)::sl2dGFP]* | This study |
| SGA70 | *N<sub>2</sub>; gaaEx070 [Punc-25::TWK-40(GF)::sl2dGFP]* | This study |
| SGA64 | *twk-40(hp834); gaaEx064 [Ptwk-40::TWK-40(GF)::sl2dGFP]* | This study |
| SGA76 | *twk-40(hp834); gaaEx076 [Punc-47::TWK-40(GF)::sl2dGFP]* | This study |
| SGA77 | *twk-40(hp834); gaaEx077 [Punc-25::TWK-40(GF)::sl2dGFP]* | This study |
| SGA89 | *twk-40(bln336); hpIs468; gaaEx089 [Ptwk-40::TWK-40(WT)::sl2dwCherry]* | This study |
| SGA98 | *twk-40(bln336); hpIs468; gaaEx098 [Punc-47::TWK-40(WT)::sl2dwCherry]* | This study |
| SGA95 | *twk-40(bln336); hpIs468; gaaEx095 [Punc-25::TWK-40(WT)::sl2dwCherry]* | This study |
| SGA90 | *twk-40(hp834); hpIs468; gaaEx090 [Ptwk-40::TWK-40(WT)::sl2dwCherry]* | This study |
| SGA93 | *twk-40(hp834); hpIs468; gaaEx093 [Punc-47::TWK-40(WT)::sl2dwCherry]* | This study |
| SGA96 | *twk-40(hp834); hpIs468; gaaEx096 [Punc-25::TWK-40(WT)::sl2dwCherry]* | This study |
| SGA86 | *hpIs468; gaaEx086 [Ptwk-40::TWK-40(GF)::sl2dwCherry]* | This study |
| SGA94 | *hpIs468; gaaEx094 [Punc-47::TWK-40(GF)::sl2dwCherry]* | This study |
| SGA91 | *hpIs468; gaaEx091 [Punc-25::TWK-40(GF)::sl2dwCherry]* | This study |
| SGA280 | *N2; gaaEx0112 [Punc-47::GFP::3'UTR; Pmyo-2::GFP]* | This study |
| SGA87 | *twk-40(hp834); hpIs468; gaaEx087 [Ptwk-40::TWK-40(GF)::sl2dwCherry]* | This study |
| SGA97 | *twk-40(hp834); hpIs468; gaaEx097 [Punc-47::TWK-40(GF)::sl2dwCherry]* | This study |
| SGA92 | *twk-40(hp834); hpIs468; gaaEx092 [Punc-25::TWK-40(GF)::sl2dwCherry]* | This study |
| SGA112 | *twk-40(bln336); hpIs582; gaaEx0111 [Ptwk-40::TWK-40(WT)::sl2dwCherry]* | This study |
| SGA290 | *twk-40(bln336); hpIs582; gaaEx0118 [Punc-47::TWK-40(WT)::sl2dwCherry]* | This study |
| SGA111 | *twk-40(bln336); hpIs582; gaaEx0110 [Punc-25::TWK-40(WT)::sl2dwCherry]* | This study |
| SGA297 | *twk-40(bln336); hpIs582; gaaEx0122 [Pexp-1::TWK-40(WT)::sl2dwCherry]* | This study |
| SGA106 | *twk-40(hp834); hpIs582; gaaEx0106 [Ptwk-40::TWK-40(WT)::sl2dwCherry]* | This study |
| SGA107 | *twk-40(hp834); hpIs582; gaaEx0107 [Punc-47::TWK-40(WT)::sl2dwCherry]* | This study |
| SGA291 | *twk-40(hp834); hpIs582; gaaEx0119 [Punc-25::TWK-40(WT)::sl2dwCherry]* | This study |
| SGA298 | *twk-40(hp834); hpIs582; gaaEx0123 [Pexp-1::TWK-40(WT)::sl2dwCherry]* | This study |
| SGA108 | *twk-40(hp834); hpIs582; gaaEx0108 [Ptwk-40::TWK-40(GF)::sl2dwCherry]* | This study |
| SGA105 | *twk-40(hp834); hpIs582; gaaEx0105 [Punc-47::TWK-40(GF)::sl2dwCherry]* | This study |
| SGA338 | *twk-40(hp834); hpIs582; gaaEx0145 [Punc-25::TWK-40(GF)::sl2dwCherry]* | This study |
| SGA299 | *twk-40(hp834); hpIs582; gaaEx0124 [Pexp-1::TWK-40(GF)::sl2dwCherry]* | This study |
| SGA996 | *twk-40(gf); hpIs468; gaaEx1251 [Pflp-10::TWK-40(WT)::sl2eTagRFP]* | This study |
| SGA997 | *hpIs468; gaaEx1252 [Pflp-10::TWK-40(GF)::sl2eTagRFP]* | This study |
| SGA998 | *twk-40(lf); hpIs468; gaaEx1253 [Pflp-10::TWK-40(GF)::sl2eTagRFP]* | This study |
| SGA999 | *twk-40(lf); hpIs468; gaaEx1254 [Pflp-10::TWK-40(WT)::sl2eTagRFP]* | This study |
| SGA1000 | *twk-40(gf); hpIs582; gaaEx1255 [Pflp-10::TWK-40(WT)::sl2eTagRFP]* | This study |
| SGA1001 | *twk-40(lf); hpIs582; gaaEx1256 [Pflp-10::TWK-40(WT)::sl2eTagRFP]* | This study |
| SGA1002 | *twk-40(lf); hpIs582; gaaEx1257 [Pflp-10::TWK-40(GF)::sl2eTagRFP]* | This study |

**Supplementary Table 2.**

| Plasmids | Detail | Source |
| --- | --- | --- |
| SG331 | *Ptwk-40(slot1)* | This study |
| SG42 | *Punc-47(slot1)* | This study |
| SG22 | *Punc-25(slot1)* | This study |
| SG324 | *Pexp-1(slot1)* | This study |
| SG137 | *Pges-1(slot1)* | This study |
| SG332 | *TWK-40(WT)(slot2)* | This study |
| SG333 | *TWK-40(GF)(slot2)* | This study |
| SG62 | *GFP(slot2)* | This study |
| SG64 | *RFP(slot2)* | This study |
| SG44 | *sl2dGFP(slot3)* | This study |
| SG144 | *sl2dwCherry(slot3)* | This study |
| SG334 | *Ptwk-40::GFP::3'UTR* | This study |
| SG335 | *Punc-47::RFP::3'UTR* | This study |
| SG336 | *Ptwk-40::TWK-40(WT)::sl2dGFP* | This study |
| SG337 | *Punc-47::TWK-40(WT)::sl2dGFP* | This study |
| SG338 | *Punc-25::TWK-40(WT)::sl2dGFP* | This study |
| SG339 | *Pges-1::TWK-40(WT)::sl2dGFP* | This study |
| SG340 | *Ptwk-40::TWK-40(GF)::sl2dGFP* | This study |
| SG341 | *Punc-47::TWK-40(GF)::sl2dGFP* | This study |
| SG342 | *Punc-25::TWK-40(GF)::sl2dGFP* | This study |
| SG343 | *Ptwk-40::TWK-40(WT)::sl2dwCherry* | This study |
| SG344 | *Punc-47::TWK-40(WT)::sl2dwCherry* | This study |
| SG345 | *Punc-25::TWK-40(WT)::sl2dwCherry* | This study |
| SG346 | *Pexp-1::TWK-40(WT)::sl2dwCherry* | This study |
| SG347 | *Ptwk-40::TWK-40(GF)::sl2dwCherry* | This study |
| SG348 | *Punc-47::TWK-40(GF)::sl2dwCherry* | This study |
| SG349 | *Punc-25::TWK-40(GF)::sl2dwCherry* | This study |
| SG350 | *Pexp-1::TWK-40(GF)::sl2dwCherry* | This study |
| SG632 | *Pflp-10::TWK-40(WT)::sl2eTagRFP* | This study |
| SG633 | *Pflp-10::TWK-40(GF)::sl2eTagRFP* | This study |

